# Transcriptional analysis defines TCR and cytokine-stimulated MAIT cells as rapid polyfunctional effector T cells that can coordinate the immune response

**DOI:** 10.1101/600189

**Authors:** Rajesh Lamichhane, Marion Schneider, Sara M. de la Harpe, Thomas W. R. Harrop, Rachel F. Hannaway, Peter Dearden, Joanna R. Kirman, Joel D. A. Tyndall, Andrea J. Vernall, James E. Ussher

## Abstract

MAIT cells are an abundant innate-like T cell population which can be activated via either their T cell receptor (TCR), which recognizes MR1-bound pyrimidine antigens derived from microbial riboflavin biosynthesis, or via cytokines, such as IL-12 and IL-18. *In vivo*, these two modes of activation may act in concert or independently depending upon the nature of the microbial or inflammatory stimuli. It is unknown, however, how the MAIT cell response differs to the different modes of activation. Here, we define the transcriptional and effector responses of human MAIT cells to TCR and cytokine stimulation. We report that MAIT cells rapidly respond to TCR stimulation through the production of multiple effector cytokines and chemokines, alteration of their cytotoxic granule content and transcription factor expression, and upregulation of co-stimulatory proteins CD40L and 4-1BB. In contrast, cytokine-mediated activation is slower and results in more limited production of cytokines, chemokines, and co-stimulatory proteins; differences in granule content and transcription factor expression are also evident. Therefore, we propose that in infections by riboflavin-synthesizing bacteria, MAIT cells play a key early role in effecting and coordinating the immune response, while in the absence of TCR stimulation (e.g. viral infection) their role is likely to differ.

## Introduction

Mucosal associated invariant T (MAIT) cells are innate-like T cells, which are abundant in blood, mucosal surfaces, and liver in humans (Martin et al., 2009). MAIT cells express a semi-invariant T cell receptor (TCR) consisting of Vα7.2-Jα12/20/33 combined with limited Vβ diversity (Gherardin et al., 2016). MAIT cells were originally discovered as CD4^−^CD8beta^−^ T cells (Tilloy et al., 1999), and are mostly CD8alpha^+^, expressing an effector memory phenotype (Leeansyah, Loh, Nixon, & Sandberg, 2014). They also share some features of natural killer (NK) cells, notably high expression of the C-type lectin CD161 (Billerbeck et al., 2010; Dusseaux et al., 2011; Kurioka, Klenerman, & Willberg, 2018).

Unlike conventional T cells, whose TCRs recognize peptides loaded on MHC molecules, the MAIT cell TCR recognizes bacteria-derived vitamin B precursor metabolites, recently identified as pyrimidine derivatives of 5-amino-6-D-ribitylaminouracil (5-A-RU), loaded on the non-polymorphic MHC class Ib related molecule 1 (MR1) (Corbett et al., 2014; S. Huang et al., 2005; Kjer-Nielsen et al., 2012; Treiner et al., 2003). Consistent with this, many riboflavin producing bacteria, but not riboflavin deficient bacteria, have been shown to stimulate MAIT cells in a MR1 dependent fashion (Gold et al., 2010; Le Bourhis et al., 2013). Various studies have reported killing of bacterially infected cells by MAIT cells *in vitro* (Kurioka et al., 2015; Le Bourhis et al., 2013) and their importance in controlling bacterial infection in mice and humans, further establishing MAIT cells as important players in anti-bacterial immunity (Chua et al., 2012; Georgel, Radosavljevic, Macquin, & Bahram, 2011; Grimaldi et al., 2014; Le Bourhis et al., 2010; Meierovics, Yankelevich, & Cowley, 2013; Smith, Hill, Bell, & Reid, 2014; Wang et al., 2018).

MAIT cells express high amounts of the interleukin-18 receptor (IL-18R) and IL-12R, and are able to respond to cytokine signals independent of their TCR (Ussher et al., 2014). This was initially shown with the combination of IL-12 and IL-18 (Jo et al., 2014; Ussher et al., 2014). IL-15 can also activate MAIT cells, but IL-15-induced MAIT cell activation is IL-18 dependent (Sattler, Dang-Heine, Reinke, & Babel, 2015). MAIT cells can be activated in a TCR independent manner by antigen presenting cells treated with bacteria (Ussher et al., 2016; Wallington, Williams, Staples, & Wilkinson, 2018) and by virally infected cells (Loh et al., 2016; van Wilgenburg et al., 2016). Additionally, MAIT cells are activated in both acute and chronic viral infections, and in patients with systemic lupus erythematous (Chiba et al., 2017; van Wilgenburg et al., 2016). Cytokine-mediated activation is important *in vivo*, as demonstrated by a recent study in mice which showed IL-18-dependent MAIT cell accumulation in lungs and MAIT cell-specific protection against challenge with a lethal dose of influenza virus (Wilgenburg et al., 2018). The *in vivo* response to bacteria includes not only a TCR-dependent but also a TCR-independent component driven by cytokines. The synergy between TCR signals and cytokine mediated signals to achieve sustained activation of MAIT cells has also been observed *in vitro* (Slichter et al., 2016; Turtle et al., 2011; Ussher et al., 2014).

In recent years, the MAIT cell phenotype has been extensively explored (Dias, Leeansyah, & Sandberg, 2017; Kurioka et al., 2017), however, the full array of effector functions of MAIT cells upon activation is yet to be completely defined. Importantly, it remains unclear whether functions driven through the TCR are distinct from those driven in a TCR-independent manner. In this study, we have explored the full range of effector functions of TCR versus cytokine activated human MAIT cells. Focusing on early events following triggering, we found marked differences in the production of cytokines, chemokines, and cytotoxic granule contents, and the expression of transcription factors in MAIT cells depending upon the mode of activation. These distinct early functional responses may reflect different roles for MAIT cells in response to diverse infectious threats and potentially in response to commensal organisms.

## 2 Materials and Methods

### 2.1 Cells and bacteria

Peripheral blood was collected from healthy donors with approval from the University of Otago Human Ethics Committee; written informed consent was obtained from all donors. PBMCs were elutriated by density gradient centrifugation using Lymphoprep^TM^ (Axis-Shield PoC AS, Oslo, Norway) and stored in liquid nitrogen. Each experiment was performed after overnight resting of thawed and washed PBMCs in RPMI 1640 supplemented with L-glutamine (Life Technologies Corporation, Carlsbad, USA), 10% fetal calf serum (Gibco, New Zealand and Sigma-Aldrich, St. Louis, USA), and Penicillin-Streptomycin (Sigma-Aldrich) (now onwards referred to as R10). For some experiments, Vα7.2^+^ cells were isolated from PBMCs by labelling with anti-Vα7.2-PE antibody (Clone; 3C10, BioLegend, San Diego, USA) followed by separation with anti-PE magnetic beads using MS columns (both from Miltenyi Biotec, Bergisch Gladbach, Germany). THP1 monocytic cells were maintained in R10.

*Escherichia coli* HB101 was cultured overnight in Luria-Bertani (LB) broth, washed with PBS (Oxoid Ltd, Basingstoke, UK) and fixed with 2% formaldehyde for 20 minutes at 4 °C. After washing, fixed bacteria were resuspended in PBS and quantified with 123count eBeads^TM^ (eBioscience, San Diego, USA) by flow cytometry using a FACSCanto II (BD biosciences, San Jose, USA).

### 2.3 MAIT cell activation

Viable PBMCs were counted by a Neubauer hemocytometer excluding dead cells using Trypan Blue (Invitrogen, Carlsbad, USA) and 10^6^ PBMCs/200 µL were seeded in U-bottom 96 well-plates. Formaldehyde-fixed *Escherichia coli*, 5-A-RU (synthesized in School of Pharmacy, University of Otago and stored in single use aliquots at −80 °C, refer to Supplementary Methods for details), or the combination of 50 ng/mL IL-12 (Miltenyi Biotec) and 50 ng/mL IL-18 (R & D Systems, Minneapolis, USA), were added at the start of each experiment. Other than in the time course experiments, PBMCs were treated with 5-A-RU or *E. coli* for 6 or 24 hours or IL-12+IL-18 for 24 hours. Some experiments were performed with 5-A-RU/MG (5-A-RU and methyl glyoxal (MG, from Sigma-Aldrich) mixed at a molar ratio of 1:50 immediately prior to use). For assessing the role of KDM6B, 10^5^ CD8^+^ T cells were treated overnight with 5 µM GSK-J4 (Sigma-Aldrich) and co-cultured with 100 bacteria per cell (BpC) *E. coli* fed 10^5^ THP1 monocytes. MR1 mediated activation of MAIT cells was blocked with 2.5 µg/mL anti-MR1 antibody (clone 26.5, BioLegend, San Diego, USA). When assessing cytokine production and cytotoxic granule content by flow cytometry, brefeldin A (BioLegend) was added at 5 µg/mL for final 4 hours of culture.

### 2.4 Flow cytometry

Cells were stained for surface markers, followed by fixing in 2% paraformaldehyde (except in sorting experiments), and treated with permeabilization buffer (BioLegend) for intracellular staining of cytokines and cytotoxic molecules or Foxp3/transcription factor buffer set (eBioscience) for transcription factors. MAIT cells were identified as CD8^+^ T-cells expressing semi-invariant TCR Vα7.2 and high levels of CD161 (Dusseaux et al., 2011); the gating strategy is shown in S. Figure 1. In some experiments, double negative and CD4^+^ MAIT cells were also examined. In all flow cytometry experiments, dead cells were excluded by live/dead fixable near IR (Invitrogen) staining. Antibodies used were: anti-CD3 PE-Cy7 (UCHT1, BioLegend), anti-CD3 BV510 (OKT3, BioLegend), anti-CD8 eFluor450 (RPA-18, eBioscience), anti-TCR Vα7.2 PE or PE-Cy7 or AF700 (3C10, BioLegend), anti-CD161 APC (191B8, Miltenyi Biotech), anti-CD161 BV605 (HP-3G10, BioLegend), anti-TCR γδ BV510 (B1, BioLegend), anti-CCR7 PerCP-Cy5.5 (G043H7, BioLegend), anti-TNFα FITC (Mab11, BioLegend), anti-IFNγ PerCP-Cy5.5 (4S.B, BioLegend), anti-IL-26 PE (510414, R & D Systems), anti-CD56 APC-Cy7, anti-CD107a PE (H4A3, BioLegend), anti-granzyme A PE (CB9, BioLegend), anti-granzyme B FITC (QA16AO2, BioLegend), anti-perforin PerCP-Cy5.5 (B-D48, BioLegend), anti-FasL PE (NOK-1, BioLegend), anti-CD40L FITC (24-31, BioLegend), anti-4-1BB PE (4B4-1, BioLegend), anti-RORγt PE (Q21-559, BD Biosciences, San Jose, CA, USA), anti-PLZF AF647 (R17-809, BD Biosciences), anti-T-bet PECy7 (4B10, BioLegend), anti-EOMES eFluor660 (WD1928, eBiosciences) and anti-Blimp1 PECF594 (6D3, BD Biosciences). Samples were acquired on a FACS CantoII, LSR Fortessa, or in sorting experiments a FACSAria^TM^ II (all BD Biosciences), and analyzed by FlowJo^TM^ V10 (TreeStar, Ashland, USA).

### 2.5 Quantitative reverse transcriptase-polymerase chain reaction (RT-PCR)

A total of 10^7^ human PBMCs were treated with formaldehyde-fixed *E. coli* (10 BpC) or 5-A-RU (5 µM) or IL-12+IL-18 (50 ng/mL each) for different durations or were left untreated. Cells were stained with fluorescently-labelled antibodies and MAIT cells (CD3^+^CD8^+^TCRγδ^−^CCR7^−^ CD161^++^Vα7.2^+^ cells) were isolated by fluorescence activated cell sorting on a FACSAria^TM^ II. RNA from sorted MAIT cells was isolated using the Nucleospin RNA isolation kit (Macherey-Nagel, Düren, Germany) as per the manufacturer’s instructions and stored at −80 °C. cDNA was synthesized using the Superscript IV system (Invitrogen) using oligo-dT primers. Real time PCR for gene expression was performed using the KAPA SYBR FAST mastermix on a ViiA7 or QS6 Real-Time system (Applied Biosystems, Foster City, USA). Previously published primers were used: TNFα (forward, AGCCTCTTCTCCTTCCTGATCGTG; reverse, GGCTGATTAGAGAGAGGTCCCTGG) (Giribaldi et al., 2010); IFNγ (forward, TGACCAGAGCATCCAAAAGA; reverse, CTCTTCGACCTCGAAACAGC) (Munk et al., 2011); IL-17A (forward, AACCGATCCACCTCACCTT; reverse, GGCACTTTGCCTCCCAGAT) (Guenova et al., 2015); IL-22 (forward, GCAGGCTTGACAAGTCCAACT; reverse, GCCTCCTTAGCCAGCATGAA) (Guenova et al., 2015); GAPDH (forward, CAACAGCGACACCCACTCCT; reverse, CACCCTGTTGCTGTAGCCAAA) (Maess, Sendelbach, & Lorkowski, 2010).

### 2.6 RNA Sequencing

A total of 10^7^ human PBMCs per donor (n=7) were treated with formaldehyde-fixed *E. coli* (10 BpC) or 5-A-RU (5 µM) for 6 hours or IL-12+IL-18 (50 ng/mL each) for 24 hours and MAIT cells were isolated by fluorescence activated cell sorting RNA was sent to the Molecular Genetics Facility at the Liggins Institute, Auckland, New Zealand for library preparation and sequencing. Libraries were prepared with the Ion AmpliSeq™ Transcriptome Human Gene Expression Kit and were sequenced on an Ion Proton. Read count data files were generated with Torrent Suite^TM^ software. Downstream analysis for differentially expressed genes (DEG) were performed using DESeq2 package in R-Bioconductor (Love, Huber, & Anders, 2014). Log_2_ fold change and adjusted p-value (Padj) were arbitrarily set at 1 (fold change 2) and 0.05 respectively as cut-offs during DEG analysis. Principal component analysis (PCA) plot was generated in DESeq2. Venn-diagrams were created from DEGs in an online tool developed by Bioinformatics & Evolutionary genomics, VIB/UGent (http://bioinformatics.psb.ugent.be/webtools/Venn/). Normalized gene expression values of total and custom lists of DEGs against all stimuli were used to generate heat-maps on Heatmapper (Babicki et al., 2016) (http://www.heatmapper.ca/expression/) employing the average linkage clustering and Spearman rank correlation distance measurement methods. Gene set enrichment analysis was performed on GSEA software version 3.0 (Mootha et al., 2003; Subramanian et al., 2005) using normalized gene expression values of all genes from 28 samples as the expression database and compared either with the curated gene set databases; KEGG or Reactome, provided in the software/website or the tissue repair gene dataset obtained from the publication by Linehan et al (Linehan et al., 2018).

### 2.7 LEGENDplex immunoassays

A total of 10^5^ column purified Vα7.2^+^ MAIT cells per donor were co-cultured with 10^5^ THP1 monocytes and stimulated with either formaldehyde-fixed *E. coli* (100 BpC) or 5-A-RU (5 µM)± anti-MR1 antibody for 6 hours; for controls, THP1 monocytes were treated with *E. coli* or 5-A-RU or were co-cultured with Vα7.2^+^ cells without any treatment. For cytokine stimulation, 10^5^ Vα7.2^+^ MAIT cells were treated with IL-12+IL-18 for 24 hours; untreated Vα7.2^+^ cells were used as a control. Cytokines, chemokines, and cytotoxic molecules in the supernatant were quantified by LEGENDplex bead-based immunoassay (BioLegend) following the manufacturer’s instructions using three multiplex panels: Human Proinflammatory Chemokine (mix and match 7-plex subpanel for CCL3, CCL4, CCL20, IL-8, CXCL9, CXCL10, and CXCL11), Human CD8/NK (mix and match 11-plex subpanel for IL-2, IL-6, IL-10, IL-17A, TNFα, IFNγ, sFasL, granzyme A, granzyme B, perforin, and granulysin), and Human custom panel (IL-1α, IL-1β, IL-17F, IL-21, IL-22, IL-23, and CD40L).

### 2.8 Neutrophil migration assay

Neutrophils were freshly isolated from peripheral blood by a two-step method of dextran (Thermofischer Scientific) sedimentation of erythrocytes followed by Lymphoprep^TM^ gradient centrifugation and removal of low-density PBMCs (Kuhns, Long Priel, Chu, & Zarember, 2015). Residual erythrocytes from the high-density neutrophil fraction were removed by lysis with water. 3×10^5^ neutrophils in 200µL R10 were added into the upper compartment of a transwell system incorporating polycarbonate membrane insert (5µm pore size, Corning), with 500 µL diluted supernatant (1:2.5 in R10) of TCR-or cytokine-stimulated MAIT cells (obtained as mentioned in LEGENDplex immunoassays) in the lower compartment; 5 ng/mL IL-8 (BD Biosciences) was used as positive control for neutrophil migration. After incubation at 37 °C for 1 hour, the transwell insert was carefully removed and the number of neutrophils in the lower compartment were counted using a hemocytometer.

### 2.9 Statistical analysis

Data were analyzed using GraphPad Prism V7.03. Statistical significance was assessed with a one-way repeated measures ANOVA with Sidak’s multiple comparison tests for multiple groups and paired two tailed t-tests for two groups. Data were normalized, or log transformed in some cases when not normally distributed. Mean ± standard error of mean (S.E.M) are shown throughout.

## 3. Results

### 3.1 Timing of maximal activation and cytokine profile of MAIT cells differ between TCR and cytokine stimulation

First, we treated human PBMCs with *E. coli* or IL-12+IL-18 and confirmed robust MAIT cell activation via TCR-dependent and -independent mechanisms (Figure 1A and B). The response to *E. coli* at 6 hours was completely MR1-dependent while at 24 hours it was only partially MR1-dependent (Figure 1A). The time course of the responses to both stimuli was then assessed at both the protein and the mRNA levels (Figure 1C-D and F-G). MAIT cells rapidly responded to *E. coli* with the transcription of multiple inflammatory cytokines (tumor necrosis factor α (TNFα), interferon γ (IFNγ), IL-17A and IL-22) compared to the untreated control; the maximum response was observed at 4-6 hours (Figure 1C). Consistently, production of TNFα and IFNγ peaked at ~ 6 hours (Figure 1D). While MAIT cells were also robustly activated with IL-12+IL-18, the response was dominated by the production of IFNγ at both mRNA and protein levels and peaked at 24 and 20 hours respectively (Figure 1E and F). Therefore, MAIT cells respond rapidly to TCR stimulation and are polyfunctional, producing multiple pro-inflammatory cytokines, whereas cytokine driven activation is slower and is dominated by IFNγ production.

**Figure 1:**
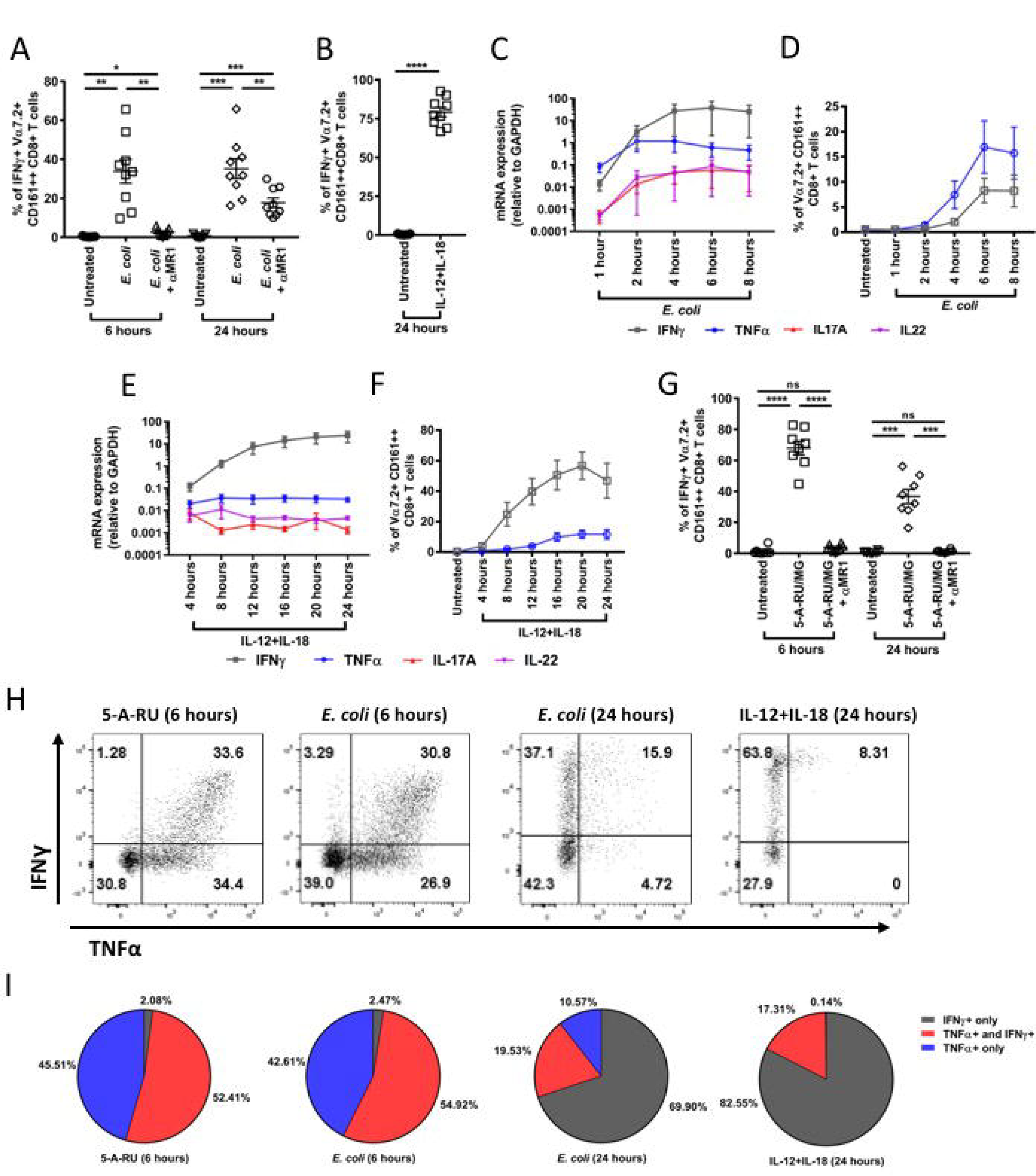
T cell receptor and cytokine-activated MAIT cells differ in timing of maximal activation and cytokine profile. **(A, B and G)** PBMCs were treated with either (A) 10 BpC *E. coli ±* anti-MR1 antibody for 6 and 24 hours or (B) 50 ng/mL each of IL-12+IL-18 for 24 hours or (G) 10 nM 5-A-RU/MG *±* anti-MR1 antibody for 6 and 24 hours and IFNγ production by MAIT cells was measured by flow cytometry. Each biological replicate and mean ± S.E.M are shown and are pooled from two independent experiments (n=9 for A and B, n=8 for G). Repetitive measures one-way ANOVA with Sidak multiple comparison (A and G) and two tailed paired t-test (B) were used for assessing statistical significance. *p<0.05, **p<0.01, ***p<0.001,****p<0.0001, ns = non-significant. **(C-F)** PBMCs were stimulated for different durations by either *E. coli* (10 BpC for C and 2 BpC for D) or by IL-12+IL-18 (50 ng/mL each) (E and F) and expression of TNFα, IFNγ, IL-17A and IL-22 in flow-sorted MAIT cells were measured by real-time RT-PCR (n=3 for C and n=4 for E) or the percentage of MAIT cells producing TNFα or IFNγ were determined by flow cytometry (D and F; n=6). Data are presented as mean ± S.E.M and are pooled from two independent experiments. (**H and I**) Proportion of MAIT cells producing TNFα or IFNγ or both were assessed following treatment of PBMCs with either 10 BpC *E. coli* or 5 µM 5-A-RU for 6 hours or 50 ng/mL each of IL-12+IL-18 for 24 hours; (E) representative FACS plots and (F) pie chart to show proportion of TNFα and IFNγ producing MAIT cells among the activated MAIT cell population against each stimulus (n=9-11). Data are pooled from two independent experiments.

Since bacteria provide signals in addition to the MAIT cell activating ligand (Ussher et al., 2016), we assessed the response to the MR1 precursor ligand, 5-A-RU, as a pure TCR signal. MAIT cell activation with 5-A-RU at both 6 and 24 hours was predominantly TCR mediated (Figure 1G). Interestingly, MAIT cells stimulated with *E. coli* or 5-A-RU for 6 hours showed similar cytokine profiles, which were markedly different from those obtained by IL-12+IL-18 stimulation for 24 hours; with TCR activation, TNFα mono-producers and TNFα/IFNγ double-producers dominated while very few IFNγ mono-producing cells were observed, whereas with cytokine stimulation IFNγ mono-producers dominated with very few TNFα/IFNγ double-producers seen (Figure 1H and I). These similar cytokine profiles of *E. coli* and 5-A-RU activated MAIT cells and the complete dependence of IFNγ production on MR1-TCR interaction during early *E. coli* treatment (Figure 1A) strongly suggests that the early bacterial activation of MAIT cells is purely TCR dependent. In contrast, *E. coli* treatment for 24 hours, which involved both TCR-dependent and-independent mechanisms (Figure 1A), elicited a cytokine profile similar to that induced by IL-12+IL-18 stimulation, although TNFα mono-producers were only seen with *E. coli* (Figure 1H and I). These results suggest that the two MAIT cell activating pathways not only differ in the timing of activation, but also the resulting cytokine production profiles.

### 3.2 Unique transcriptional profiles of TCR-and cytokine-stimulated MAIT cells

To further investigate the differences observed with the two modes of activation, transcriptomic profiles of MAIT cells activated via TCR with either *E. coli* or 5-A-RU for 6 hours or via cytokine receptors with IL-12+IL-18 for 24 hours were generated by RNA sequencing; untreated MAIT cells were included for comparison. First, we compared the response of MAIT cells to different treatments by principal component analysis. We observed good separation of activated MAIT cells from non-activated MAIT cells across the first two principal components that together accounted for 62% of total variability in the dataset (Figure 2A). MAIT cells activated by *E. coli* or 5-A-RU formed overlapping clusters that were separated from cytokine treated MAIT cells; this difference was largely driven by principal component 1 (PC1). We also observed some similarity between IL-12+IL-18 and *E. coli* treatments in PC2.

**Figure 2:**
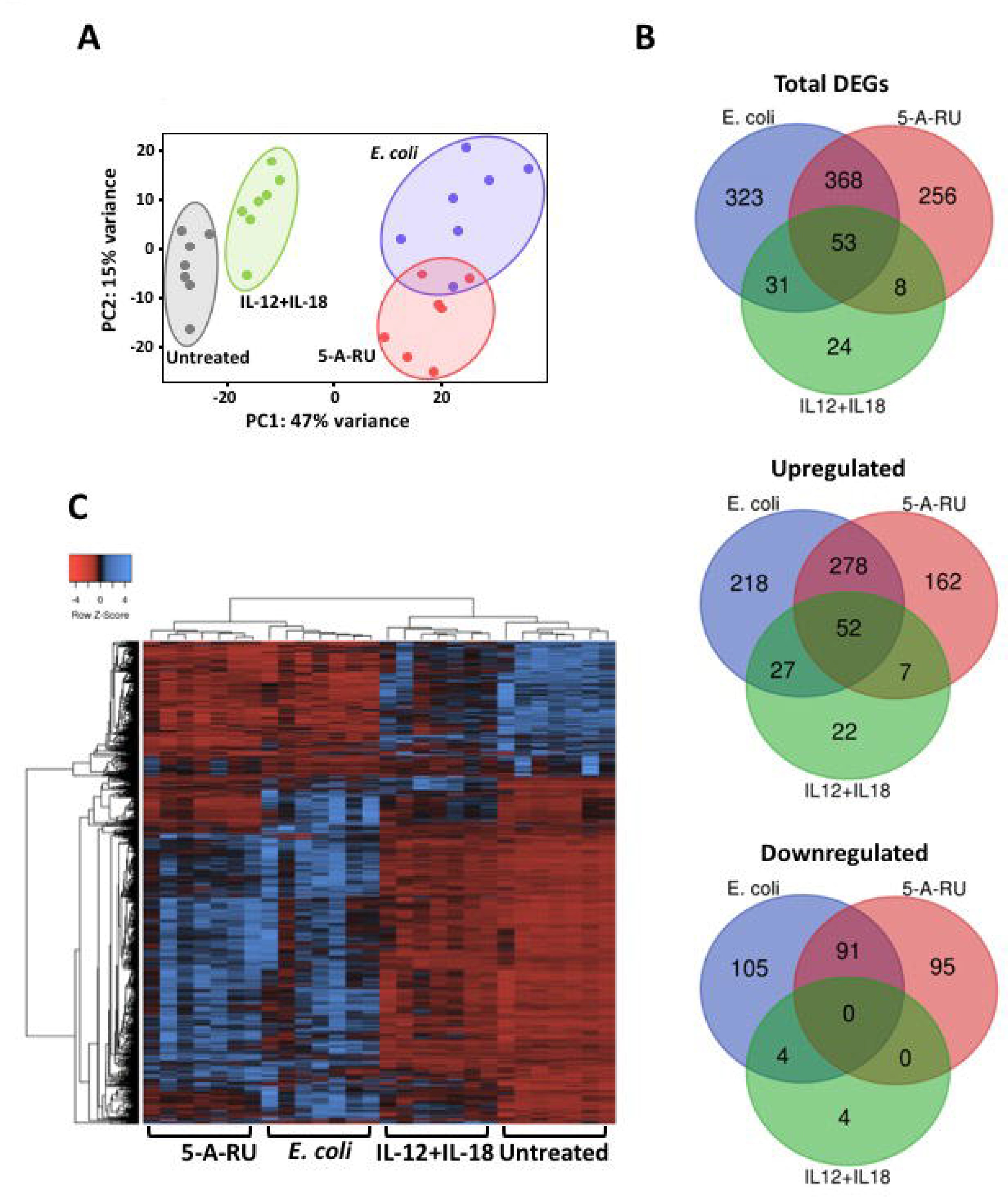
T cell receptor and cytokine-activated MAIT cells have distinct transcriptional profiles. (**A**) Principal component analysis of MAIT cell transcriptome following stimulation with different stimuli; PBMCs were treated with 10 BpC *E. coli* or 5 µM 5-A-RU for 6 hours or 50 ng/mL IL-12+IL-18 for 24 hours, then MAIT cells were flow-sorted for RNA-sequencing. Principle components (PC) 1 and 2 are shown; each dot represents a sample and are color coded for ease of visualization. (**B**) Venn-diagram showing shared and unique DEGs (fold change >2 and Padj <0.05) in TCR-and cytokine-stimulated MAIT cells; total, upregulated and downregulated DEGs with each treatment compared to untreated control were separately analyzed. (**C**) Heat map and dendrogram of the normalized expression of total DEGs (1063) with all treatments compared to untreated control.

Next, differentially expressed genes (DEGs) in activated MAIT cells with respect to rested MAIT cells were analyzed. Substantial transcriptional changes in MAIT cells were observed with the treatments; DEGs were more abundant with either *E. coli* (775) or 5-A-RU (685) treatment than with IL-12+IL-18 (116). Interestingly, more than a quarter of DEGs were downregulated with both *E. coli* and 5-A-RU but very few (8) with IL-12+IL-18 (Figure 2B). A significant number of DEGs (53; which was 45% of total DEGs with cytokine stimulation) were shared in all three treatments, suggesting a core response of activated MAIT cells regardless of the mode of activation. This was evident with GSEA pathway analysis with pathways such as NK cell mediated cytotoxicity (S. Figure 2A), cytokine-cytokine receptor interaction, T cell receptor signaling pathway, apoptosis, NOD-like receptor signaling, and JAK/STAT signaling enriched with *E. coli*, 5-A-RU, and IL-12+IL-18. Of note, IL-12+IL-18 stimulated MAIT cells shared more DEGs (84; 72% of total) with *E. coli* treated MAIT cells but less (61; 52% of total) with 5-A-RU, suggesting bacteria and innate cytokines activate a similar response which cannot be triggered by the MR1 ligand alone. GSEA confirmed that many innate signaling pathways that were enriched with *E. coli* when compared to 5-A-RU were also enriched with IL-12+IL-18 (S. Table 1, S. Figure 2B). Transcriptomic signatures associated with TCR triggering could also be identified. A novel tissue repair signature proposed to be associated with TCR triggering of MAIT cells (Leng et al., 2018) was enriched in MAIT cells following *E. coli* or 5-A-RU stimulation but not with IL-12+IL-18 (S. Figure 2C).

A heat-map illustrating the normalized expressions of total DEGs with all stimuli also showed marked differences in the response of MAIT cells at transcriptomic level; activation with *E. coli* or 5-A-RU had more pronounced effects on gene expression and were clustered separately from the cluster of untreated and cytokine treated MAIT cells (Figure 2C). DEGs with each treatment are listed in Supplementary Data. Taken together, our data suggest MAIT cells are differentially regulated by TCR and cytokine stimuli and possess a unique transcriptomic profile from that of resting MAIT cells.

### 3.3 MAIT cells activated via TCR produce more inflammatory cytokines than IL-12+IL-18-stimulated MAIT cells

We next investigated the effect of *E. coli*, 5-A-RU, or IL-12+IL-18 on the production of inflammatory cytokines by MAIT cells. With all stimuli, increased expression of TNFα and IFNγ genes was seen; nevertheless, higher TNFα expression was achieved with TCR stimuli and maximum IFNγ expression with IL-12+IL-18 (Figure 3A). Consistent with this, more TNFα was detected in the culture supernatant following stimulation with *E. coli* than with IL-12+IL-18; more TNFα was produced in response to *E. coli* than to 5-A-RU (>10-fold) (Figure 3B). In contrast, more than 10-fold higher concentration of IFNγ was detected upon IL-12+IL-18 stimulation for 24 hours compared to stimulation with *E. coli* for 6 hours; ~100-fold more IFNγ was produced in response to *E. coli* than to 5-A-RU (Figure 3B). MR1 blocking significantly inhibited TNFα and IFNγ production by both *E. coli* and 5-A-RU treated MAIT cells (Figure 3B). TCR stimulation also triggered significant upregulation of expression of several other inflammatory genes, notably IL1-α, IL-1β, IL-2, IL-6, IL-10, IL-17A/F, IL-21, IL-22, and GM-CSF, while with IL-12+IL-18 stimulation significant upregulation of IL-10 and IL-17A/F was seen, but to a much lesser degree than with TCR mediated activation (Figure 3A, 3C and S. Figure 4). Of the cytokines assayed in the supernatant, we detected MR1-dependent production of IL-1β, IL-2, IL-6, and IL-10 in response to *E. coli* but not 5-A-RU, however, only IL-1β and IL-2 were significant (Figure 3D). Low level production of IL-1β, IL-6, and IL-10 was detected in response to IL-12+IL-18 (Figure 3D). Therefore, MAIT cells produce more pro-inflammatory cytokines in response to TCR stimulation than in response to IL-12+IL-18; the response to *E. coli* is stronger than to 5-A-RU.

**Figure 3:**
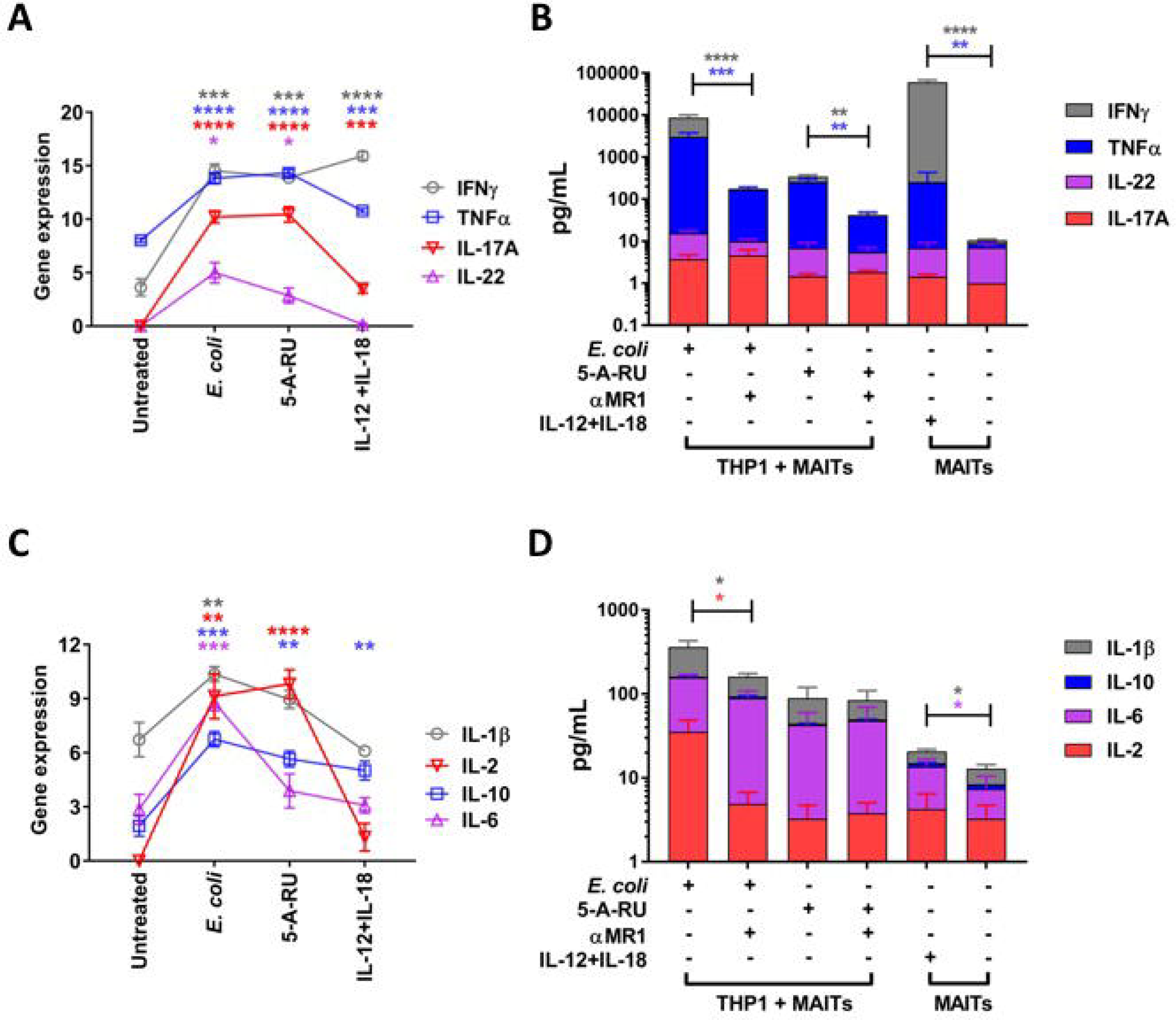
TCR stimulation results in production of more inflammatory cytokines than IL-12+IL-18 stimulation. (**A and C**) Log_2_ transformed gene counts of TNFα, IFNγ, IL-17A, IL-22, IL-1β, IL-2, IL-6, and IL-10 in response to different stimuli; PBMCs were treated with 10 BpC *E. coli* or 5 µM 5-A-RU for 6 hours or 50 ng/mL IL-12+IL-18 for 24 hours, then MAIT cells were flow-sorted for RNA sequencing. Data are presented as mean ± S.E.M (n=7). Repetitive measures one-way ANOVA with Sidak multiple comparison tests were used for assessing statistical significance. *p<0.05, **p<0.01, ***p<0.001, ****p<0.0001. (**B and D**) Cytokines (TNFα, IFNγ, IL-17A, IL-22, IL-1β, IL-2, IL-6, and IL-10) were quantified by LEGENDplex^TM^ in either culture supernatant from co-cultures of purified Vα7.2^+^ cells and THP1 monocytes (1:1 ratio) stimulated with 100 BpC *E. coli* or 5 µM 5-A-RU for 6 hours ± anti-MR1, or culture supernatant of purified Vα7.2^+^ cells alone or after IL-12+IL-18 treatment for 24 hours. Data are presented as mean ± S.E.M and are pooled from two independent experiments (n=5). Paired t-test were performed on log-transformed data for assessing statistical significance. *p<0.05, **p<0.01, ***p<0.001, ****p<0.0001.

Interestingly, cytokine stimulation, and to a lesser degree TCR stimulation, triggered significant upregulation of IL-26 gene expression, a proinflammatory cytokine of the IL-10 superfamily (S. Figure 5A). Likewise, cytokine activated MAIT cells expressed IL-26 at protein level, the highest amongst CD8^+^ T cells (S. Figure 5B and C), but early TCR stimulation failed to induce detectable IL-26 production (S. Figure 5D and E). However, IL-26 production was detectable upon late TCR activation by *E. coli* and 5-A-RU/MG and was significantly inhibited by MR1 blocking (S. Figure 5D and E). At 24 hours, both *E. coli* and 5-A-RU/MG treated MAIT cells produced a small amount of IL-26 in an MR1-independent fashion (S. Figure 5D and E). IL-26 production was also observed in NK cells (identified as CD3^−^CD161^+^Vα7.2^−^ cells, ~80% of which were CD56^+^) following stimulation with IL-12+IL-18 or treatment with *E. coli* for 24 hours; the response to *E. coli* was independent of MR1 (S. Figure 5F-H). Overall, this suggests that MAIT cells are the main source of IL-26 among CD8^+^ T cells in response to IL-12+IL-18 and that they also produce it in response to TCR stimulation, but production is delayed.

### 3.4 Cytotoxic profile of MAIT cells is different between the two modes of activation

MAIT cells can kill bacterially infected cells (Le Bourhis et al., 2013), which is mostly degranulation dependent (Kurioka et al., 2015). We asked whether degranulation and the cytotoxic granule content of MAIT cell differs with the mode of activation. First, we assessed MAIT cell degranulation by assessing surface expression of CD107a. CD107a surface expression rapidly increased with both TCR stimuli and with IL-12+IL-18, suggesting both modes of activation are equally potent at triggering degranulation in MAIT cells (Figure 4A). Consistent with this, *E. coli*-mediated CD107a expression at 6 hours was significantly reduced with MR1 blocking, whereas only partial inhibition was observed at 24 hours, confirming the previous finding of both TCR-dependent and -independent late degranulation in MAIT cells (Figure 4A) (Kurioka et al., 2015).

**Figure 4:**
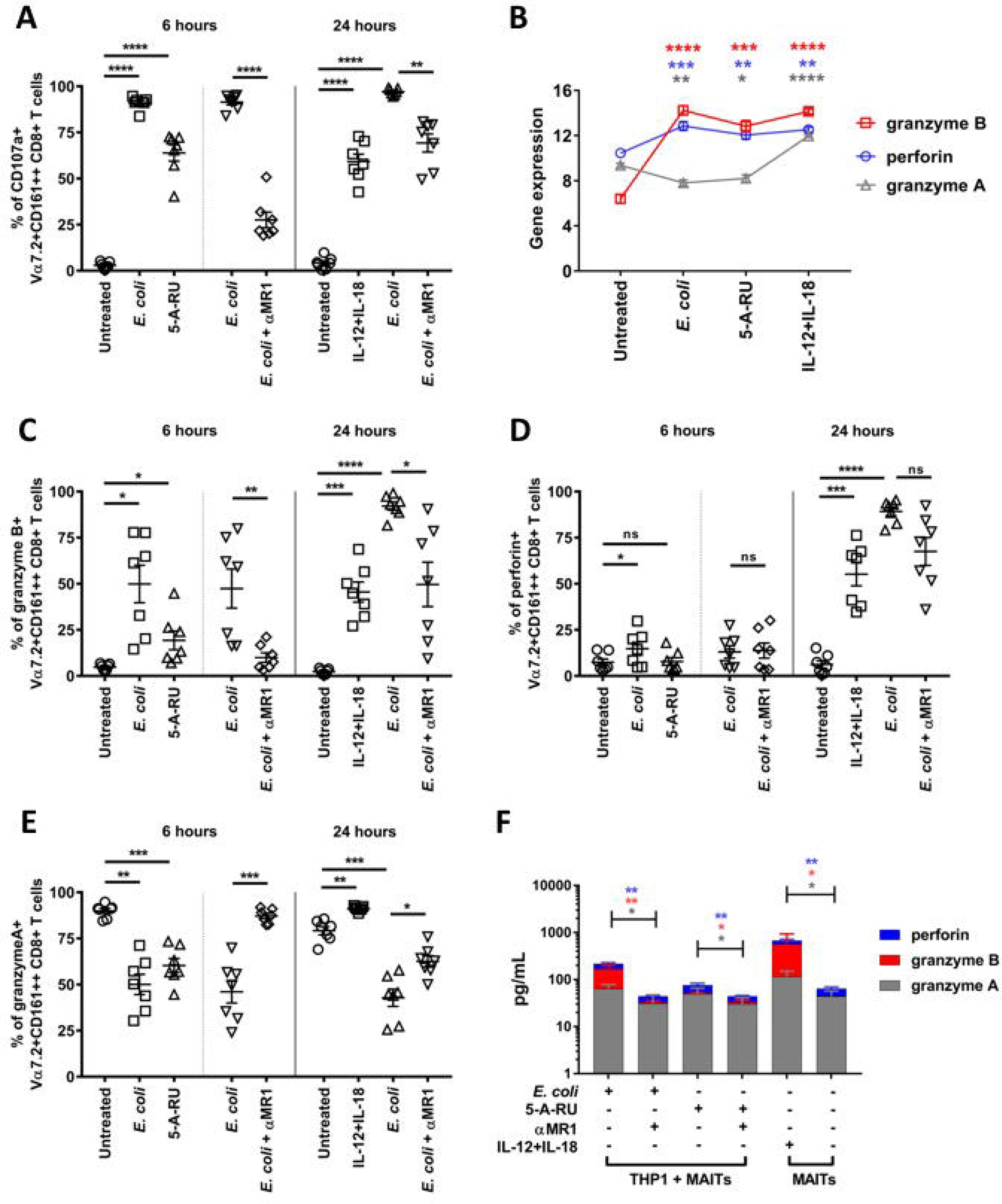
Cytotoxic granule content of MAIT cells differs with the mode of activation. (**A, C-E**) PBMCs were stimulated with either 10 BpC *E. coli* or 5 µM 5-A-RU for 6 hours or 50 ng/mL IL-12+IL-18 or 10 BpC *E. coli* for 24 hours and the percentage of CD107a (A), granzyme B (C), perforin (D), and granzyme A (E) expressing MAIT cells were measured by flow cytometry; with *E. coli* stimulation, the effect of anti-MR1was assessed. Each biological replicate and mean ± S.E.M are shown and are pooled from two independent experiments (n=7). Repetitive measures one-way ANOVA with Sidak multiple comparison and two tailed paired t-tests were used for statistical analysis. *p<0.05, **p<0.01, ***p<0.001, ****p<0.0001, ns = non-significant. (**B**) Comparison of log_2_ transformed gene counts of granzyme B, perforin and granzyme A in response to different stimuli; PBMCs were treated with 10 BpC *E. coli* or 5 µM 5-A-RU for 6 hours or 50 ng/mL IL-12+IL-18 for 24 hours, then MAIT cells were flow-sorted for RNA sequencing. Data are presented as mean ± S.E.M (n=7). Repetitive measures one-way ANOVA with Sidak multiple comparison tests were used for assessing statistical significance. *p<0.05, **p<0.01, ***p<0.001, ****p<0.0001. (**F**) Cytotoxic molecules were quantified by LEGENDplex^TM^ in either culture supernatant from co-cultures of purified Vα7.2^+^ cells and THP1 monocytes (1:1 ratio) stimulated with 100 BpC *E. coli* or 5 µM 5-A-RU for 6 hours ± anti-MR1, or culture supernatant of purified Vα7.2^+^ cells alone or after IL-12+IL-18 treatment for 24 hours. Data are represented as mean ± S.E.M and are pooled from two independent experiments (n=5). Paired t-test were performed on log transformed data for assessing statistical significance. *p<0.05, **p<0.01.

Next, we compared cytotoxic granule content of MAIT cells upon activation. Activated MAIT cells upregulated both granzyme B and perforin gene expression irrespective of the mode of activation (Figure 4B). Likewise, the frequency of MAIT cells expressing granzyme B significantly increased upon both early TCR and cytokine stimulation (Figure 4C). While the frequency of MAIT cells expressing perforin increased with *E. coli* and cytokines, it did not with 5-A-RU (Figure 4D). Nevertheless, 5-A-RU treatment for 24 hours induced significant granzyme B and perforin expression which was completely TCR mediated (S. Figure 6A and B). MR1 blockade abolished *E. coli*-induced granzyme B production completely at 6 hours and partially at 24 hours but had no effect on perforin production at 6 hours and minimal effect at 24 hours (Figure 4C and D). Interestingly, *E. coli* or 5-A-RU treatment triggered granzyme A gene downregulation in MAIT cells but IL-12+IL-18 treatment resulted in upregulation (Figure 4B). This difference in granzyme A expression was also evident at protein level; both early and late TCR-mediated activation reduced granzyme A whereas IL-12+IL-18 significantly increased expression (Figure 4E). *E. coli*-mediated granzyme A downregulation was reversed completely with MR1 blocking at 6 hours and partially at 24 hours (Figure 4E). Granzyme A, granzyme B, and perforin were detected in the culture supernatant of both TCR and cytokine stimulated MAIT cells; however, more granzyme A was detected in the supernatant of cytokine treated MAIT cells (Figure 4F). Differences were also evident in gene expression of other granzymes; granzymes K and M followed a similar trend to granzyme A while granzyme H followed that of granzyme B except it was highest with IL-12+IL-18 stimulation (S. Figure 6C). Overall, these data suggest that MAIT cells rapidly degranulate upon activation, however, granule content differs by the mode and timing of activation.

Cytotoxic T cells can also kill target cells through expression of FasL. Both TCR and cytokine activated MAIT cells rapidly upregulated FasL gene expression (S. Figure 6D). Although, FasL could not be detected on the surface of early activated MAIT cells, soluble FasL was detected in the culture supernatant of early TCR activated MAIT cells in a MR1-dependent fashion (S. Figure 6E and F). Late *E. coli* activation significantly increased FasL expression on the surface and was only partially reduced upon MR1 blocking (S. Figure 6E). IL-12+IL-18 activation enhanced both FasL surface expression and sFasL release into the media (S. Figure 6E and F) suggesting both activation pathways contribute to FasL/sFasL expression.

### 3.5 TCR and cytokine activated MAIT cells express different profiles of transcription factors

MAIT cells are characterized by high expression of the transcription factors RAR-related orphan receptor γt (RORγt), promyelocytic leukemia zinc finger protein (PLZF) and eomesodermin (EOMES), which make them unique from other CD4^+^ and CD8^+^ T cells (Figure 5A) (Dias et al., 2017; Leeansyah et al., 2015). Expression of these transcription factors also varies among different MAIT cell subsets based on CD4 or CD8 co-receptor expression, resulting in heterogeneity in phenotype and function (S. Figure 7A) (Kurioka et al., 2017). Therefore, we assessed the expression of various transcription factors in MAIT cells to investigate whether the transcription factor profile changes with the mode of activation. Interestingly, compared to resting MAIT cells, the expression of RORγt and PLZF significantly increased with TCR activation but not with IL-12+IL-18 (Figure 5B and C). We observed corresponding changes in RORγt expression at the protein level but little change in PLZF expression were observed (Figure 5A-C). Upon activation, MAIT cells expressed high T-bet regardless of the stimuli, both at RNA and protein levels (Figure 5A, C and D). Interestingly, with TCR stimulation an increase in MAIT cells expressing both RORγt and T-bet or RORγt alone were seen, whereas following IL-12+IL-18 treatment an increase in MAIT cells expressing T-bet alone was observed (Figure 5G). Expression of the Blimp1 gene went up with both TCR stimuli and with cytokines, however, an increase at the protein level was only seen with *E. coli* and cytokines (Figure 5E). EOMES expression, both at the mRNA and protein levels, was significantly reduced by both TCR signals but was unchanged with cytokines (Figure 5F). Similar changes in the transcription factors in response to TCR signals and IL-12+IL-18 were evident in double negative and CD4^+^ MAIT cell subsets (S. Figure 7B-F). Taken together, the profile of transcription factor expression in MAIT cells markedly changes upon activation and is different in TCR and cytokine stimulated MAIT cells.

**Figure 5:**
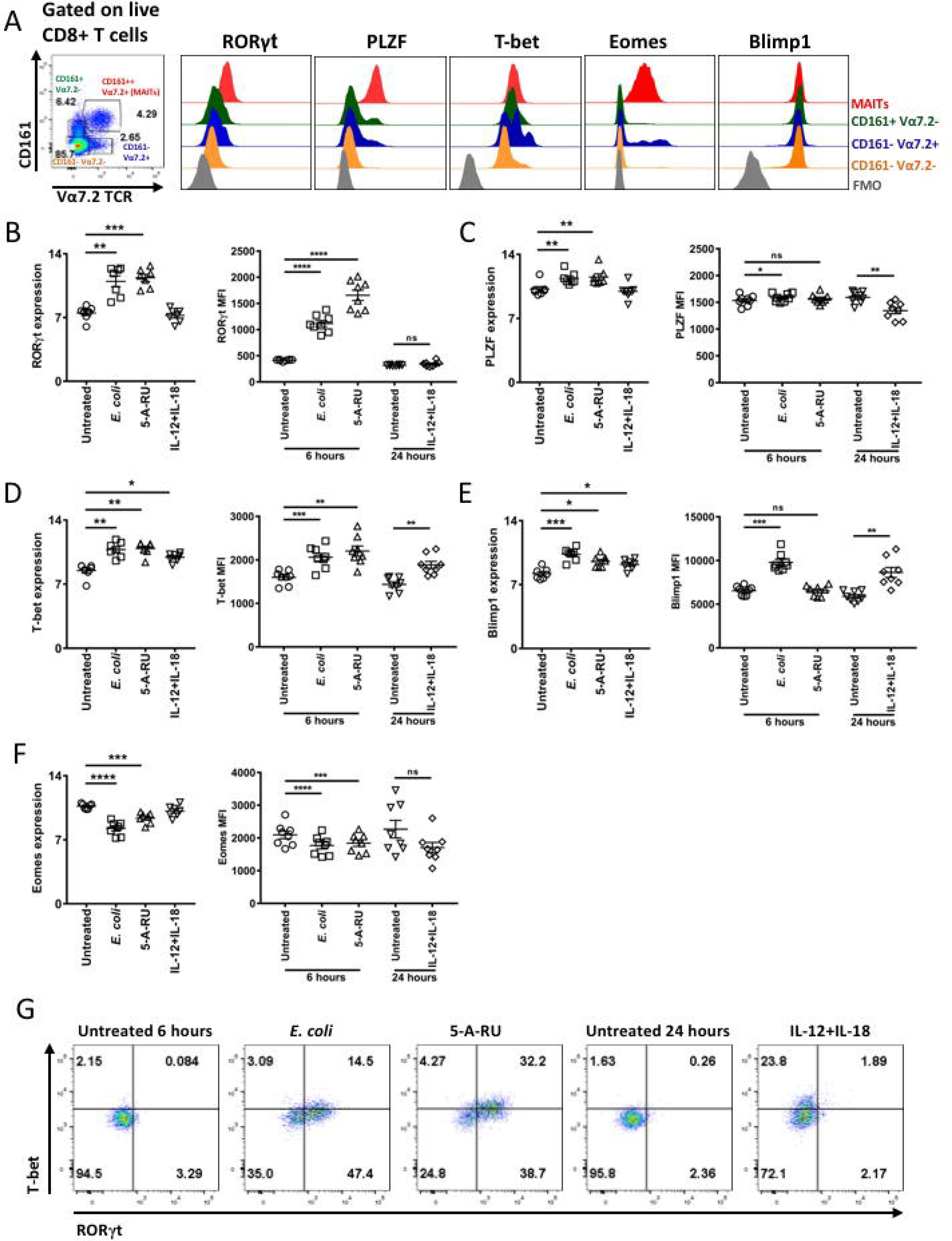
MAIT cells express a unique transcription factor profile which changes upon activation. (**A**) Expression of different transcription factors were compared among MAIT cells and other non-MAIT CD8^+^ T cells from PBMCs; representative histograms and the gating strategy for MAIT and non-MAIT cells is shown. An FMO (fluorescent minus one) control for each transcription factor was included. (**B-F**) Human PBMCs were treated with 10 BpC *E. coli* or 5 µM 5-A-RU for 6 hours or IL-12+IL-18 (50 ng/mL each) for 24 hours and expression of different transcription factors were compared, both at RNA level (log_2_ transformed gene counts) on isolated MAIT cells, and at protein level directly in PBMCs by measuring MFI on MAIT cells; (B) RORγt, (C) PLZF, (D) T-bet, (E) Blimp1, and (F) EOMES. Each biological replicate and mean ± S.E.M are shown and are pooled from two independent experiments (n=7 for RNA and n=8 for protein). Repetitive measures one-way ANOVA with Sidak multiple comparison were performed for statistical analysis. *p<0.05, **p<0.01, ***p<0.001, ****p<0.0001, ns = non-significant. (**G**) Upregulation of RORγt alone or T-bet alone or both on MAIT cells upon TCR-or cytokine-activation were compared; representative plots are shown.

### 3.6 TCR stimulation of MAIT cells results in chemokine production

Production of chemokines by MAIT cells upon activation has previously been shown (Lepore et al., 2014; Slichter et al., 2016; Turtle et al., 2011). Next, we examined the expression of various chemokines by MAIT cells in response to different stimuli. Significant expression of CCL3 (ligand for CCR1, CCR4 and CCR5), CCL4 (ligand for CCR5), and CCL20 (ligand for CCR6) was seen with all treatments but was significantly higher in MAIT cells activated with *E. coli* and 5-A-RU than with IL-12+IL-18 (Figure 6A). Consistent with this, CCL3, CCL4, and CCL20 production was detected in culture supernatant with *E. coli* or 5-A-RU treatment and was reduced by approximately 10-fold upon MR1 blocking, confirming production was MAIT cell specific (Figure 6B). Cytokine stimulation also triggered significant CCL3 and CCL4 production, albeit at lower levels (Figure 6B). We also observed enhanced CXCL9, CXCL10, and CXCL11 gene expression, all ligands for CXCR3, with 5-A-RU and IL-12+IL-18 treatment, whereas *E. coli* treatment resulted in a lesser, non-significant enhancement (Figure 6C). In contrast, at the protein level, *E. coli* stimulated maximum CXCL9, CXCL10, and CXCL11 production by MAIT cells, followed by treatment with 5-A-RU and IL-12+IL-18 (Figure 6D); MR1 blocking completely abolished production of CXCL9, CXCL10, and CXCL11 by *E. coli* and 5-A-RU treated MAIT cells (Figure 6D).

**Figure 6:**
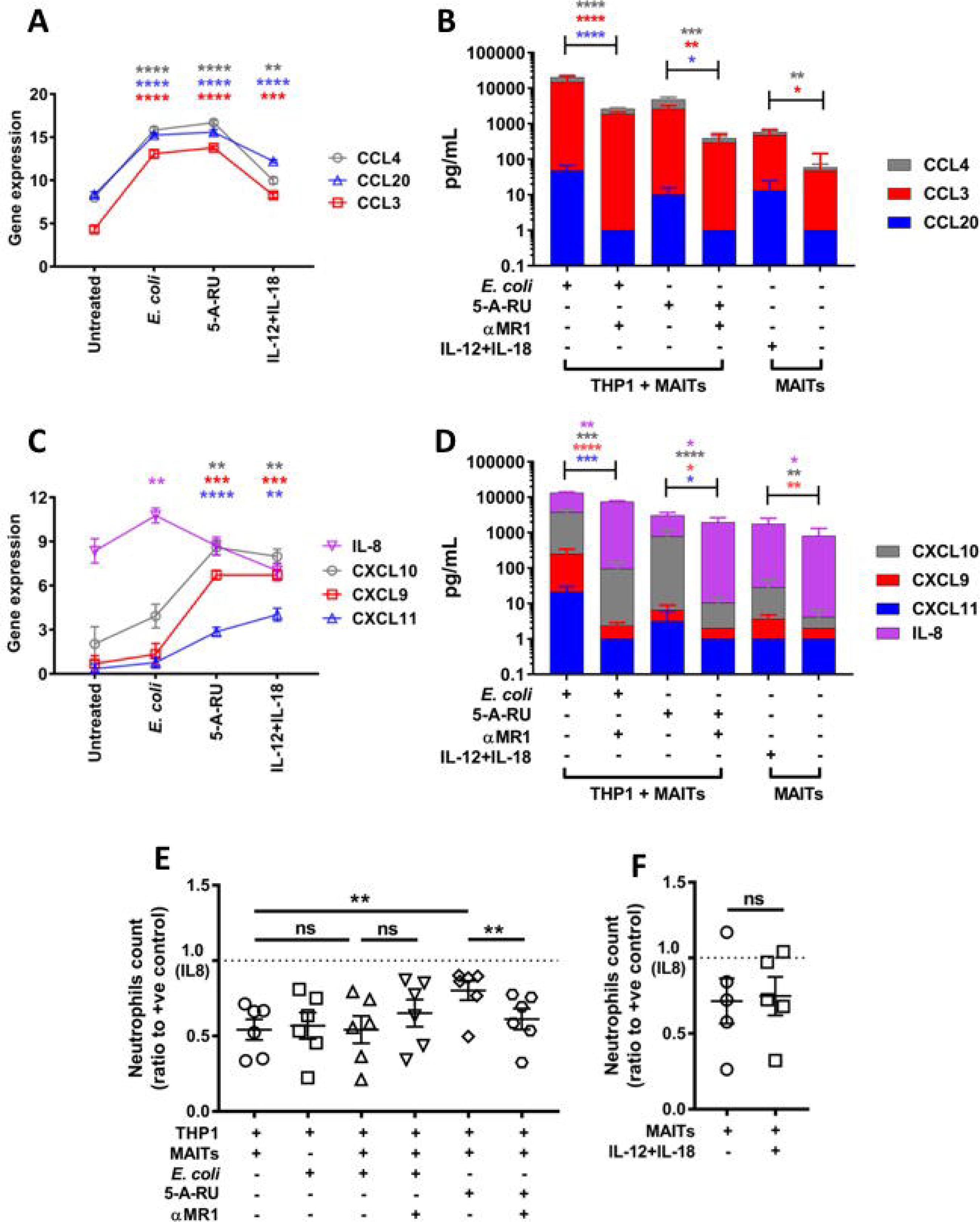
Robust chemokine production in MAIT cells following TCR activation. (A and C) Comparison of log_2_ transformed gene counts of CCL3, CCL4, CCL20, CXCL9, CXCL10, CCL20, and IL-8 in response to different stimuli; PBMCs were treated with 10 BpC *E. coli* or 5 µM 5-A-RU for 6 hours or 50 ng/mL IL-12+IL-18 for 24 hours, then MAIT cells were flow-sorted for RNA sequencing. Data are presented as mean ± S.E.M (n=7). Repetitive measures one-way ANOVA with Sidak multiple comparison tests were used for assessing statistical significance. **p<0.01, ***p<0.001, ****p<0.0001. (**B and D**) Chemokines (CCL3, CCL4, CCL20, CXCL9, CXCL10, CCL20, and IL-8) were quantified by LEGENDplex^TM^ in either culture supernatant from co-cultures of purified Vα7.2^+^ cells and THP1 monocytes (1:1 ratio) stimulated with 100 BpC *E. coli* or 5 µM 5-A-RU for 6 hours ± anti-MR1, or culture supernatant of purified Vα7.2^+^ cells alone or after IL-12+IL-18 treatment for 24 hours. Data are presented as mean ± S.E.M and are pooled from two independent experiments (n=5). Paired t-test were performed on log-transformed data for assessing statistical significance. *p<0.05, **p<0.01, ***p<0.001, ****p<0.0001. **(E and F)** Migrated neutrophils in the bottom well of a transwell plate in response to supernatant of TCR-(E) or cytokine-(F) activated MAIT cells were normalized to that of the positive control (IL-8); supernatant from unactivated MAIT cells or following MR1 blockade with anti-MR1 were included as controls. Each biological replicate and mean ± S.E.M are shown; independent experiments were performed with each donor (n=6 for E and n=5 for F). Repetitive measures one-way ANOVA with Sidak multiple comparison (E) and two-tailed paired t-test (F) were used. **p<0.01, ns = non-significant.

Additionally, *E. coli* but not 5-A-RU or IL-12+IL-18, resulted in upregulation of IL-8 (CXCL8) gene expression (Figure 6C). Consistent with this, IL-8 production was highest in supernatant from *E. coli* treated MAIT cells and was significantly blocked by anti-MR1 antibody (Figure 6D). 5-A-RU and IL-12+IL-18 stimulation also caused low level but significant IL-8 production (Figure 6D). The finding of IL-8 production suggested that MAIT cells may be one of the first responders to bacterial infection and may assist in recruiting other immune cells, including neutrophils (Ribeiro, Flores, Cunha, & Ferreira, 1991). To investigate whether activated MAIT cells are able to induce neutrophil migration; supernatants from *E. coli* or 5-A-RU or IL-12+IL-18 stimulated MAIT cells were tested in a transwell migration system. Enhanced neutrophil migration was only observed in response to supernatant of 5-A-RU stimulated MAIT cells (Figure 6E-F). Blocking MR1 signaling completely abrogated the induced migration confirming that it was mediated by TCR signaling of MAIT cells (Figure 6E). CCL2 and XCL2, which both have the potential to trigger migration of neutrophils by binding to CCR2 and XCR1 respectively (Fox et al., 2015; H. Huang, Li, Cairns, Gordon, & Xiang, 2001; Talbot et al., 2015), were also upregulated with 5-A-RU stimulation compared to other treatments and may explain the increased neutrophil migration with supernatant from 5-A-RU treated MAIT cells (S. Figure 4).

Overall, MAIT cells produce multiple chemokines upon activation by both TCR-dependent and -independent mechanisms, however, chemokine profiles differ according to the mode of activation.

### 3.7 Activated MAIT cells rapidly upregulate co-stimulatory molecules

Recently, MAIT cells were associated with maturation of monocyte-derived and primary dendritic cells in a CD40L dependent manner (Salio et al., 2017). We next investigated how CD40L is controlled in MAIT cells upon activation. Both TCR and cytokine stimulated MAIT cells rapidly upregulated CD40L gene expression (Figure 7A). Unstimulated MAIT cells did not express CD40L on the surface but treatment with *E. coli* for 6 hours led to significant CD40L surface expression; in contrast little upregulation was seen with 5-A-RU (Figure 7B). MR1 blocking reduced *E. coli* mediated CD40L surface expression completely at both 6 hours and at 24 hours (Figure 7B). Significant CD40L was also seen on the surface of MAIT cells following IL-12+IL-18 stimulation (Figure 7B). Soluble CD40L could not be detected in the culture media with any treatments (data not shown).

**Figure 7:**
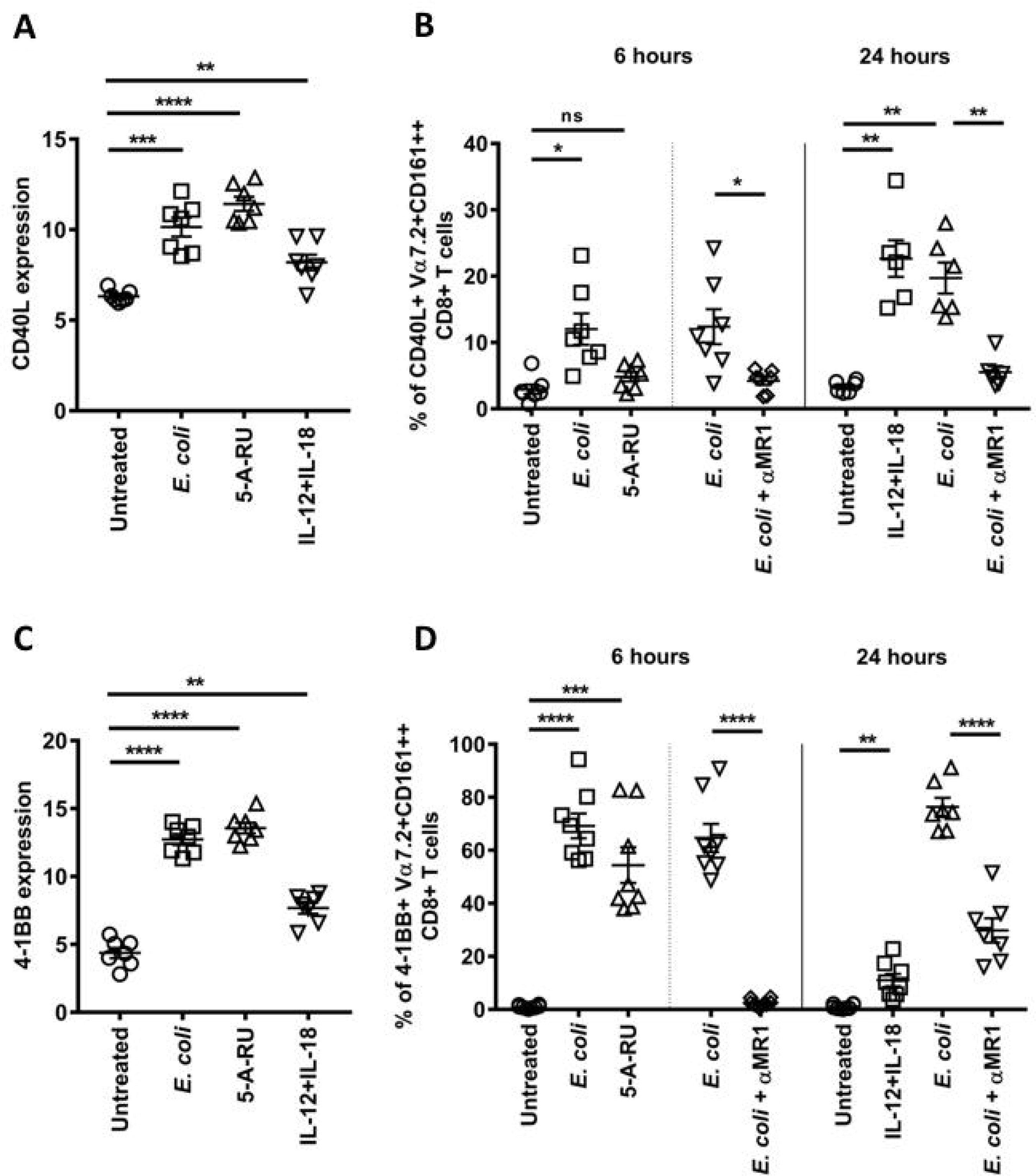
TCR and cytokine activation leads to increased expression of co-stimulatory molecules on MAIT cells. (A, C) Comparison of CD40L (A) and 4-1BB (C) log_2_ transformed gene counts in response to different stimuli; PBMCs were treated with 10 BpC *E. coli* or 5 µM 5-A-RU for 6 hours or 50 ng/mL IL-12+IL-18 for 24 hours, then MAIT cells were flow-sorted for RNA sequencing. Data are presented as mean ± S.E.M (n=7). Repetitive measures one-way ANOVA with Sidak multiple comparison tests were used for assessing statistical significance. **p<0.01, ***p<0.001, ****p<0.0001. **(B, D)** Percentage of MAITs expressing CD40L (B) or 4-1BB (D) following treatment of PBMCs with either 10 BpC *E. coli* or 5 µM 5-A-RU for 6 hours or IL-12+IL-18 or 10 BpC *E. coli* for 24 hours; in *E. coli* treatment MR1 blockade with ani-MR1 was included at 6 hours and 24 hours. Each biological replicate and mean ± S.E.M are shown and are pooled from two independent experiments (n=7 for A and C, n=6-7 for B, and n=7-8 for D). Repetitive measures one-way ANOVA with Sidak multiple comparison and two-tailed paired t-test were used for statistical analysis. *p<0.05, **p<0.01, ***p<0.001, ****p<0.0001, ns = non-significant.

We also observed increased expression of 4-1BB (TNFRSF9 or CD137), a co-stimulatory molecule on activated T cells that can trigger maturation, activation and migration of cells expressing 4-1BBL, such as dendritic cells (Lippert et al., 2008), by both TCR and cytokine activated MAIT cells, with the highest expression seen with TCR stimuli (Figure 7C). Consistent with this, high surface expression of 4-1BB was seen following early TCR activation which was completely MR1 mediated (Figure 7D). A small percentage of MAIT cells also expressed 4-1BB upon stimulation by IL-12+IL-18, suggesting cytokines alone can induce 4-1BB on MAIT cells (Figure 7D). Late TCR activation by *E. coli* resulted in high expression of 4-1BB that was only partially blocked with anti-MR1 antibody (Figure 7D). Therefore, TCR stimulation induces robust 4-1BB surface expression on MAIT cells, however cytokines alone can induce its expression, albeit to a lesser extent.

### 3.6 TCR stimulation of MAIT cells enhances expression of lysine demethylase 6B

Finally, expression of lysine demethylase 6B, KDM6B, also known as Jumonji domain containing protein 3 (JMJD3), was specifically enriched upon TCR activation (S. Figure 3A). Inhibition of KDM6B by GSK-J4 demonstrated it was important for the upregulation of CD69 and 4-1BB following stimulation with *E. coli*, but not for TNFα and IFNγ production in MAIT cells (S. Figure 3B-F).

## 4. Discussion

MAIT cells are abundant unconventional T cells that can be activated either via their TCR or by innate cytokines. Here, we have characterized their responses to these two modes of activation. TCR activation of MAIT cells resulted in rapid activation with production of multiple proinflammatory cytokines; in contrast cytokine-mediated activation was slower and less polyfunctional. Substantial differences in the cytotoxic granule content upon TCR and cytokine stimulation were also noted. These differences in effector functions by mode of activation were underpinned by changes in transcription factors expression. Production of chemokines and upregulation of co-stimulatory molecules with activation was also demonstrated and suggests a role for MAIT cells in coordinating the recruitment and activation of other immune cells.

Early TCR activation with 5-A-RU or with *E. coli* triggers higher TNFα production by MAIT cells than IL-12+IL-18. Upon stimulation with IL-12+IL-18, MAIT cells cytokine production shifts to IFNγ. This is consistent with an earlier study where human PBMCs were activated with either anti-CD3/CD28 beads or a combination of IL-12/15/18 (Slichter et al., 2016). A similar response has been reported when hepatic MAIT cells were stimulated with anti-CD3/CD28 beads (Jo et al., 2014). Predominance of IFNγ production with little TNFα production was seen when hepatic MAIT cells were treated overnight with *Pseudomonas aeruginosa*, a riboflavin synthesizing bacteria (Jo et al., 2014); this is consistent with the response we observed from blood MAIT cells following treatment with *E. coli* for 24 hours. This highlights that the early MAIT cell response to riboflavin-synthesizing bacteria differs to the late response and is consistent between blood and hepatic MAIT cells. With the observation that production of IFNγ, IL-17A and IL-22 accompany TNFα in early TCR activation, we propose that a TCR signal is critical for TNFα production and polyfunctionality of MAIT cells.

While MAIT cells stimulated via their TCR with either *E. coli* or 5-A-RU showed considerable overlap in their transcriptional and functional profiles, there were substantial differences in DEGs, which was evident in principal component 2 of the PCA analysis. Of note, cytokine stimulated MAIT cells shared more DEGs with *E.* coli than 5-A-RU, suggesting a role for innate signaling. It has now been established that bacteria furnish co-stimulatory signals, TLR agonists being most commonly studied, that result in APC activation and enhancement of MAIT cell activation (Chen et al., 2017; Ussher et al., 2016). Consistent with this, pathway analysis demonstrated similarities between *E. coli* and IL-12+IL-18 activated MAIT cells, with enrichment of innate signaling pathways including type I IFNs, pattern recognition receptors, and MYD88/TRIF. However, these signals must work in concert with TCR signaling to modulate MAIT cell activation as MR1 blockade almost completely inhibited the early MAIT cell response (IFNγ, granzyme B and chemokines) to *E. coli*.

The hypothesis that TCR triggering leads to early polyfunctionality in MAIT cells was confirmed by RNA sequencing. In addition to TNFα, IFNγ, IL-17A, and IL-22, the expression of many pro-inflammatory cytokine genes, including IL-1β, IL-6, IL-21, and GM-CSF was upregulated with TCR stimulation. However, significant production of only TNFα, IFNγ, and IL-1β could be detected in the culture supernatant of *E. coli* activated MAIT cells; GM-CSF was not assayed. These differences between gene expression and protein production could be due to the different culture systems used (PBMCs vs a co-culture of THP1 cells and Vα7.2^+^ cells) and the presence or absence of necessary additional signals, for example IL-7 which enhances IL-17A production (Tang et al., 2013). Arguing against this, however, another study (Bennett, Trivedi, Iyer, Hale, & Leung, 2017) reported significant detection of IL-2, IL-6, IL-10, IL-17A/F, IL-21, and IL-22 in the culture supernatant following overnight stimulation with *E. coli* using the same co-culture system as ours. In that study, TCR signaling was only partially responsible for the production of IL-6, IL-10, and IL-21. This may be due to the different times post-stimulation at which cytokine production was assessed. Of note, the major pro-inflammatory mediators TNFα, IL-6, and IL-1β are highly expressed at both mRNA and protein levels with *E. coli* but not with 5-A-RU or IL-12+IL-18 stimulation, suggesting that TCR activation by riboflavin synthesizing bacteria is required for some pro-inflammatory responses by MAIT cells.

A striking difference was observed in the cytotoxic granule content of MAIT cells with the two modes of activation. Perforin, granzyme A, and granzyme B were all substantially induced by IL-12+IL-18, whereas early TCR activation triggered a smaller increase in granzyme B and a decrease in granzyme A. By 24 hours, activation by *E. coli* triggered substantial granzyme B and perforin production which was partially TCR-dependent and partially -independent. Activation with the 5-A-RU for 24 hours induced significantly more granzyme B and perforin compared to activation for 6 hours; this suggests that changes in the cytotoxic granule content of MAIT cells is a time dependent process and for a strong response, either prolonged activation via the TCR, cytokines, or a combination of both is required. We and others have demonstrated granzyme A downregulation in MAIT cells with TCR signals which was an intriguing finding and demands further investigation (Kurioka et al., 2015). Conversely, granzyme A upregulation with cytokines could be of importance in the control of viral and bacterial infections as unlike granzyme B, granzyme A can potently mediate cytolysis in the presence of caspase inhibitors (Blasche et al., 2013; Zhang, Beresford, Greenberg, & Lieberman, 2001) or alternatively, may aid in the inflammatory response as discussed previously (Kurioka et al., 2015). Additionally, both modes of activation result in the upregulation of FasL on the surface of MAIT cells; FasL is involved in death receptor-mediated killing of infected cells (Varanasi, Khan, & Chervonsky, 2014).

MAIT cells may also exercise direct killing of extracellular bacteria through the release of antibacterial peptides such as IL-26 (Meller et al., 2015) and antimicrobial chemokines such as CXCL9 and CXCL10 (Margulieux, Fox, Nakamoto, & Hughes, 2016; Reid-Yu, Tuinema, Small, Xing, & Coombes, 2015). This makes MAIT cells one of many cell populations contributing to the pool of antimicrobial peptides at the site of infection/inflammation. Indirectly, MAIT cells can assist in bacterial killing by recruiting neutrophils to the site of infection, as suggested in our study, and by augmenting neutrophil survival via release of TNFα, IFNγ, and GM-CSF (Davey et al., 2014). In a positive feedback mechanism, MAIT cells could also promote DCs to produce type I interferons in an IL-26 dependent manner, which augments IFNγ release by MAIT cells in response to other cytokines (Meller et al., 2015; van Wilgenburg et al., 2016). IFNγ production in MAIT cells was recently shown to be critical for protection against *Legionella longbeachae* and influenza virus infection in mice (Wang et al., 2018; Wilgenburg et al., 2018).

Transcription factor profiling in activated MAIT cells was consistent with the observed acquired effector functions in previous studies on MAIT cells. Elevated RORγt expression was seen with TCR activation and is consistent with increased IL-17A expression (Wang et al., 2018). Both TCR and cytokine activation resulted in increased expression of T-bet, which is consistent with the substantial IFNγ production. Increased Blimp1 expression with *E. coli* and cytokines was consistent with more granzyme B production than with 5-A-RU, whereas reduced EOMES expression following early TCR stimulation was consistent with overall lower cytotoxic response of early TCR activated MAIT cells compared to IL-12+IL-18 activation (Kurioka et al., 2017; Kurioka et al., 2015). Transcription factor profiles were also assessed in other MAIT cell populations. At rest, CD4^−^CD8^−^ double negative (DN) MAIT cells expressed a similar transcription factor profile as CD8^+^ MAIT cells, consistent with the recent report where DN MAIT cells were shown to be functionally similar to CD8^+^ MAIT cells (Kurioka et al., 2017). On the other hand, CD4^+^ MAIT cells (<5% of the MAIT cell population) expressed lower levels of PLZF and EOMES, consistent with Kurioka et al, who demonstrated that they also possess lower cytotoxic potential than CD8^+^ and DN MAIT cells (Kurioka et al., 2017). Although not all changes reached statistical significance, the trends in changes of transcription factor expression upon activation were similar in all subsets, suggesting that all MAIT cells undergo similar functional changes during activation, although this was not formally assessed in this study.

KDM6B is one of the two histone demethylases (the other being KDM6A or UTX) that removes the methyl mark from H3K27, relieving the suppression of transcription and inducing an inflammatory response (Falvo, Jasenosky, Kruidenier, & Goldfeld, 2013). Murine studies have confirmed demethylated H3K27 is heavily enriched at *IL17a/IL17f* locus in Th17 cells and also controls RORγt expression (Mukasa et al., 2010). Our data suggest that upon TCR stimulation, MAIT cells upregulate KDM6B (>13 and >10-fold change with *E. coli* and 5-A-RU respectively), which could lead to the subsequent upregulation of RORγt and IL-17A expression. We also demonstrated that H3K27me3 is an important epigenetic modification during activation of MAIT cells. Conversely, IL-12 treatment of Th17 cells was previously reported to lead to a substantial increase in H3K27me3 at the *IL17a/IL17f* locus and H3K4me accumulation at the *IFNg* locus, leading to reduction of IL-17 and increased IFNγ production, consistent with our data (Mukasa et al., 2010). Thus, the mode of activation modulates the fate of MAIT cells by epigenetic remodeling.

We found that MAIT cells produce multiple chemokines upon activation which may aid in recruiting various immune subsets to the site of infection and promoting local inflammation. TCR signals induced stronger production of chemokines by MAIT cells. Previous studies have also reported substantial chemokine production by MAIT cells upon TCR activation via CD3/CD28 (Slichter et al., 2016; Turtle et al., 2011) or in a co-culture with bacteria-fed monocytes (Lepore et al., 2014; Wakao et al., 2013). However, these studies were performed with longer activation periods (16-24 hours or overnight). Here, we confirmed that MAIT cells possess this immune cell recruiting capacity as early as 6 hours after activation. We also showed that IL-12+IL-18 alone can stimulate modest chemokine production, particularly CCL3, CCL4, and CXCL10. These chemokines are involved in lymphocyte trafficking, and therefore could contribute to recruiting lymphocytes during infections with viruses or non-riboflavin-synthesizing bacteria. Moreover, a combination of cytokines and TCR signal, can work in synergy to trigger robust chemokine production by MAIT cells (Havenith et al., 2012).

In addition to this direct chemokine response, soluble mediators in the supernatant from stimulated MAIT cells have been shown to trigger the production of various chemokines, such as CCL2, IL-8, and CXCL10, by peritoneal epithelial and fibroblast cells (Liuzzi et al., 2016). MAIT cells and γδ T cells were shown to migrate to infected peritoneal tissues in response to locally elevated levels of CCL2, CCL3, CCL4, and CCL20 (Liuzzi et al., 2016). High production of CCL3, CCL4, and CCL20, along with moderate production of CXCL9 and CXCL10 by both TCR and cytokine stimulated MAIT cells were observed in our study; combined with high expression of CCR5, CCR6 and CXCR3 on MAIT cells (Dias et al., 2017; Kurioka et al., 2017), this suggests that MAIT cells, both in the presence of riboflavin-synthesizing bacteria and other inflammatory stimuli, may directly recruit other MAIT cells and/or γδ T cells (Figure 8). Furthermore, significant production and release of TNFα and IL-1β observed with bacterial stimulation can further facilitate this process by upregulating E-selectins, I-CAM and V-CAM on endothelial cells (Kim et al., 2017). Therefore, activated tissue resident MAIT cells could recruit peripheral MAIT cells, along with other proinflammatory leukocytes, to the site of infection.

**Figure 8:**
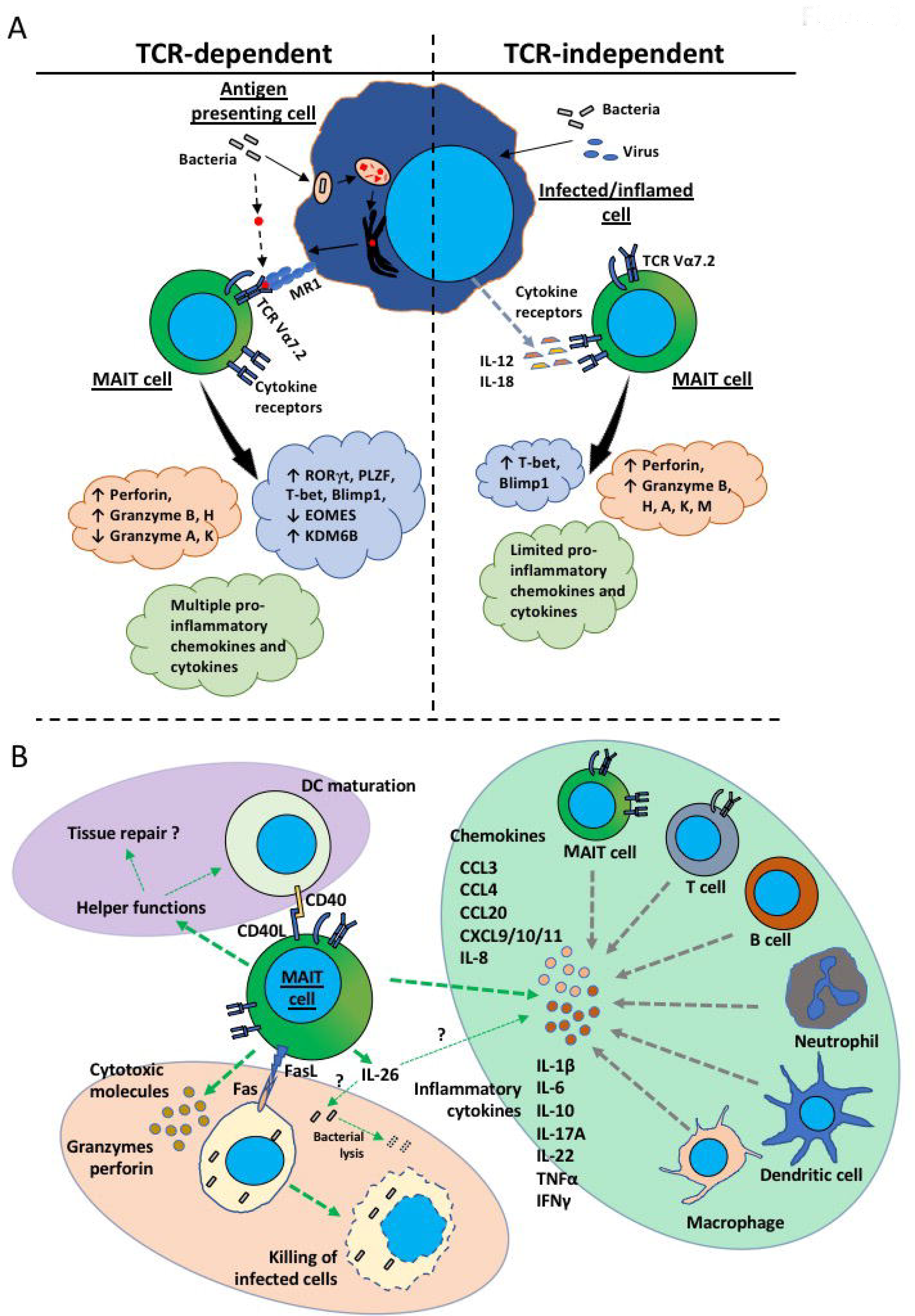
Summary of differences in TCR-and cytokine-stimulated MAIT cells (A) and role of MAIT cells in the immune response when activated via both modes during infection with riboflavin synthesizing bacteria (B).

Recent studies have demonstrated that MAIT cells do not just provide defense against invading pathogens but may have a broader role in the regulation of the immune response through the maturation of primary DCs and providing B cell help (Bennett et al., 2017; Salio et al., 2017). Both TCR and innate signals were found to upregulate helper molecules on MAIT cells. Additionally, MAIT cells were recently shown to induce local tissue remodeling (Liuzzi et al., 2016). Here, we observed specific upregulation of genes associated with tissue repair and wound healing which was recently described to be a feature of commensal responsive skin resident CD8^+^ populations in non-human primates and mice (Linehan et al., 2018). This response was more prominent with TCR activation compared to cytokine activation. Our finding was consistent with those of Leng et al., who demonstrated a similar enrichment of genes involved in tissue repair functions in TCR and TCR+cytokine but not cytokine triggered peripheral human MAIT cells (Leng et al., 2018), and Hinks et al., who reported enrichment of the tissue repair pathway in 5-OP-RU-stimulated human MAIT cells and MAIT cells from mice acutely infected with *L. longbeachae* (Hinks et al., 2018); this confirms that the tissue repair function is a unique signature of TCR activated MAIT cells. Therefore, it is now becoming clear that MAIT cells are not just inflammatory and cytolytic T cells but are also helper cells involved in tissue repair and maintaining homeostasis at the mucosal barrier and in the liver.

In this paper, we explored the response of TCR-and cytokine-stimulated MAIT cells from peripheral blood. Future investigations will shed light on whether effector function profiles of MAIT cells from different tissues also differ with the mode of activation and will enable us to understand the role of MAIT cells as a friend, foe, or merely a bystander in disease progression. The limited number of cytokines assessed in the time course studies gave us a snapshot of the timing of MAIT cell activation; other effector functions induced by a pure TCR signal should be assessed by a detailed transcriptomic analysis at later timepoints to better define the induction and persistence of particular effector functions following TCR stimulation.

In summary, our findings show that TCR and cytokines are both potent modes of triggering MAIT cell activation. MAIT cells activated via IL-12+IL-18 upregulate T-bet, perforin, granzyme B, and IFNγ and are less pro-inflammatory, consistent with a Tc1 like phenotype, while TCR-activated MAIT cells robustly and rapidly upregulate RORγt, IL-17A, TNFα, and many other pro-inflammatory cytokines and chemokines, consistent with a Tc17 phenotype (Figure 8A). This plasticity in MAIT cell phenotype and function with the mode of activation is explained by changes at transcriptomic and epigenetic levels. During bacterial infection, both modes of activation cooperate to make MAIT cells potent early responders, combining direct antimicrobial activity with the recruitment of and provision of help to other proinflammatory immune cells (Figure 8B).

## Acknowledgements

This study was funded by a grant by Health Research Council (JEU), and a University of Otago Research Grant (JDAT, AJV, JEU). We would like to thank Michelle Wilson, Flow cytometry Unit, University of Otago, for assistance with fluorescence activated cell sorting experiments, and Professor Paul Klenerman, University of Oxford, for his comments on the manuscript.

## Author contributions

RL, MS, SMdlH, and RFH performed the experiments. RL, MS, and TWRH analysed the data. RL, MS, SMdlH, JDAT, AJV, PD, JRK, and JEU designed the experiments. JEU managed the study. RL and JEU conceived the work and wrote the manuscript. All authors revised and approved the manuscript.

**S. Figure1: Flow cytometric gating strategy for phenotypic and functional analysis as well as isolation of MAIT cells.** MAIT cells were identified as CD3^+^CD8^+^CD161^++^Vα7.2^+^ cells for assessing effector functions by flow cytometry; for sorting experiments for real-time RT-PCR and RNA sequencing two additional markers (γδ-TCR and CCR7) were used to exclude γδ T cells and central memory CD8^+^ T cells respectively and MAIT cells were sorted as CD3^+^CD8^+^TCRγδ^−^CCR7^−^CD161^++^ Vα7.2^+^ cells. Number represents percentage of parent for respective population.

**S. Figure 2: Gene set enrichment analysis (GSEA) revealed shared and unique features in MAIT cells activated with *E. coli*, 5-A-RU or IL-12+IL-18**. Summary enrichment plots from GSEA analysis for (A) NK cell-mediated cytotoxicity (KEGG), (B) RIG-I like receptor signaling pathway (KEGG), and (C) tissue repair when log_2_ normalized total gene expression of *E. coli* (EC), 5-A-RU (5A), or IL-12+IL-18 (CY) stimulated MAIT cells were compared to that of untreated (UT) MAIT cells. Enrichment scores and nominal p-values are included for each analysis.

**S. Figure 3: Histone demethylase KDM6B is important for *E. coli* mediated MAIT cell activation**. (**A**) Comparison of log_2_ transformed gene counts of KDM6A and KDM6B in untreated MAIT cells (UT) and in response to different stimuli; PBMCs were treated with 10 BpC *E. coli* or 5 µM 5-A-RU for 6 hours or 50 ng/mL IL-12+IL-18 for 24 hours, then MAIT cells were flow-sorted for RNA-sequencing. Data are presented as mean ± S.E.M (n=7). Repetitive measures one-way ANOVA with Sidak multiple comparison tests was used for assessing statistical significance. **p<0.01, ****p<0.0001. (**B-F**) Column purified CD8^+^ T cells were treated overnight with KDM6B inhibitor GSK-J4 or vehicle control (DMSO) and co-cultured with THP1 monocytes. Expression of (B) CD69 and percentage of (C) 4-1BB, (D) viable, and (E) TNFα and (F) IFNγ producing MAIT cells were assessed following stimulation with 100 BpC *E. coli* for 6 hours. Each biological replicate and mean ± S.E.M are shown and are pooled from two independent experiments (n=7). Repetitive measures one-way ANOVA with Sidak multiple comparison was used for statistical analysis. **p<0.01, ****p<0.0001, ns = non-significant.

**S. Figure 4: T cell receptor and cytokine-activated MAIT cells have distinct inflammatory profiles.** Normalized expression of a custom list of genes in MAIT cells in response to different stimuli and unstimulated MAIT cells was used to generate the heat map with dendrogram using the Heatmapper online tool (Babicki et al., 2016).

**S. Figure 5: Late activation leads to increased expression of IL-26 on MAIT cells regardless of the mode of activation.** (**A**) Log_2_ transformed IL-26 count was compared in MAIT cells activated with different stimuli; PBMCs were treated with 10 BpC *E. coli* or 5 µM 5-A-RU for 6 hours or 50 ng/mL IL-12+IL-18 for 24 hours, then MAIT cells were FACS sorted for RNA-sequencing (n=7). Repetitive measures one-way ANOVA with Sidak multiple comparison tests was used for assessing statistical significance. **p<0.01, ****p<0.0001. (**B and C**) IL-26 production by MAIT cells and other non-MAIT CD8^+^ T cells was measured by intracellular staining after PBMCs were treated with 50 ng/mL IL-12+IL-18 for 24 hours. Data are presented as mean ± S.E.M and are pooled from two independent experiments (n=9). Paired t-tests were used for statistical analysis. *p<0.05, ***p<0.001. (**D and E**) PBMCs were treated with (D) 10 BpC *E. coli* or (E) 10 nM 5-A-RU/MG for 6 and 24 hours and IL-26 production by MAIT cells was assessed; anti-MR1 antibody was also added for 6 hours or 24 hours. Each biological replicate and mean ± S.E.M are shown and are pooled from two independent experiments (n=9). Repetitive measures one-way ANOVA with Sidak multiple comparison tests were used for assessing statistical significance. **p<0.01, ***p<0.001, ns = non-significant. **(F)** CD3^−^CD161^+^ cells under live lymphocytes are mainly CD56^+^ NK cells. **(G-H)** PBMCs were treated with (G) *E. coli* for 6 and 24 hours or (H) IL-12+IL-18 for 24 hours and IL-26 production by CD3^−^ CD161^+^ cells was assessed; anti-MR1 antibody was also added during *E. coli* treatments at 6 hours or 24 hours. Each biological replicate and mean ± S.E.M are shown and are pooled from two independent experiments (n=9). Repetitive measures one-way ANOVA with Sidak multiple comparison and paired t tests were performed. **p<0.01, ns = non-significant.

**S. Figure 6. Expression of cytotoxic molecules and FasL/sFasL by TCR and cytokine activated MAIT cells.** (A and B) Human PBMCs were treated with 5 µM 5-A-RU ± anti-MR1 antibody for 24 hours and percentage of MAIT cells expressing granzyme B (A) and perforin (B) were measured by flow cytometry. Each biological replicate and mean ± S.E.M are shown and are pooled from two independent experiments (n=6). Repeated measures one-way ANOVA with Sidak multiple comparison tests were used for statistical analysis. *p<0.05, ****p<0.0001, ns = non-significant**. (C and D)** Gene counts (log_2_ transformed) of (C) granzyme H, K, and M and (D) FasL with different stimuli; PBMCs were treated with 10 BpC *E. coli* or 5 µM 5-A-RU for 6 hours or 50 ng/mL IL-12+IL-18 for 24 hours, then MAIT cells were flow-sorted for RNA sequencing. Data are presented as mean ± S.E.M (n=7); for FasL, each biological replicate is shown. Repetitive measures one-way ANOVA with Sidak multiple comparison tests were used for assessing statistical significance. *p<0.05, **p<0.01, ***p<0.001, ****p<0.0001 **(E)** Percentage of FasL expressing MAIT cells after PBMCs were treated with 10 BpC *E. coli* or 5 µM 5-A-RU for 6 hours or IL-12+IL-18 (50 ng/mL each) for 24 hours; MR1 was blocked in *E. coli* stimulation for 24 hours. Each biological replicate and mean ± S.E.M are shown and are pooled from two independent experiments (n=6-7). Repetitive measures one-way ANOVA with Sidak multiple comparison and paired t tests were performed. **p<0.01, ns = non-significant. (F) Soluble FasL was quantified by LEGENDplex^TM^ in either culture supernatant from co-cultures of purified Vα7.2^+^ cells and THP1 monocytes (1:1 ratio) stimulated with 100 BpC *E. coli* or 5 µM 5-A-RU for 6 hours ± anti-MR1, or culture supernatant of purified Vα7.2^+^ cells alone or after IL-12+IL-18 treatment for 24 hours. Data are presented as mean ± S.E.M (n=5). Paired t-test were performed on log transformed data for assessing statistical significance. **p<0.01, ***p<0.001.

**S. Figure 7: Similar shift in transcription factor expression in other MAIT cell subsets during activation.** (**A**) MAIT cells were identified as CD3^+^CD161^++^Vα7.2^+^ T cells in human PBMCs and expression of different transcription factors were compared in double negative (DN; CD8^−^CD4^−^), CD4^+^, and double positive (DP; CD4^+^CD8^+^) MAIT cell subsets; representative histograms and the gating strategy for the different MAIT cell subsets are shown. An FMO (fluorescent minus one) control for each transcription factor was included. (**B-F**) Human PBMCs were treated with 10 BpC *E. coli* or 5 µM 5-A-RU for 6 hours or IL-12+IL-18 (50 ng/mL each) for 24 hours and expression of different transcription factors in different MAIT cell subsets was assessed by flow cytometry; (B) RORγt, (C) PLZF, (D) T-bet, (E) Blimp1, and (F) EOMES. Each biological replicate and mean ± S.E.M are shown and are pooled from two independent experiments (n=8). Repetitive measures one-way ANOVA with Sidak multiple comparison was performed for statistical analysis. *p<0.05, **p<0.01, ***p<0.001, ****p<0.0001, ns = non-significant.

