## Supplementary Method for "Transcriptional analysis defines TCR and cytokine-stimulated MAIT cells as rapid polyfunctional effector T cells that can coordinate the immune response"

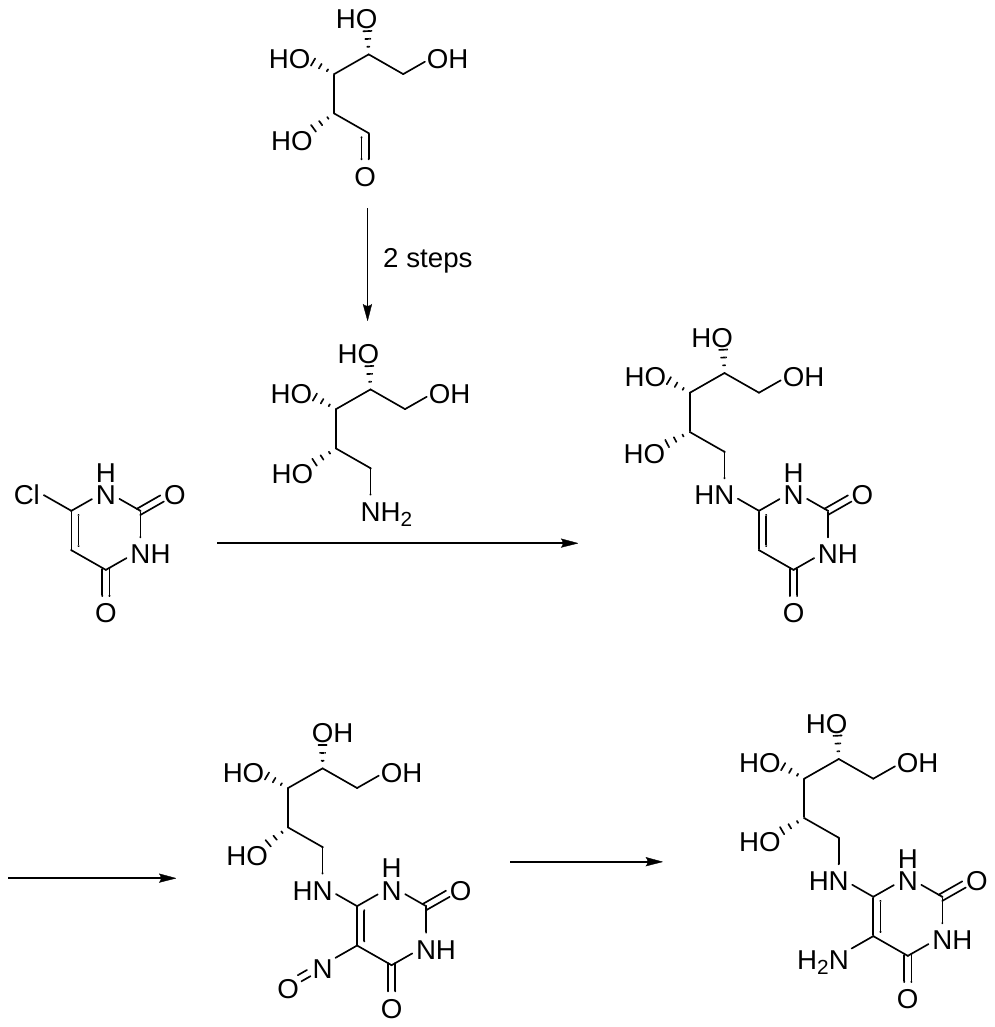

**(2R,3S,4S)‐5‐Aminopentane‐1,2,3,4‐tetrol**

Hydroxylamine hydrochloride (4.24g, 0.061 mmol) was diluted in anhydrous ethanol (24 mL) and two drops of a 1% phenolphthalein in ethanol solution were added at rt. A solution of sodium methoxide (25% wt in methanol, 13 mL) was added slowly to the suspension, upon which a white precipitate formed. Addition of the sodium methoxide was halted when the mixture stayed pink for approximately 1 min. The mixture was stirred for 3 min and filtered. The filtrate was warmed to 70°C and D-(-)-ribose ((2R,3R,4R)‐2,3,4,5‐tetrahydroxypentanal) (26.64 mmol, 4.00 g) was added in 3 batches. The mixture was stirred at 70°C until the sugar had completely dissolved, then cooled to rt. Upon cooling a precipitated formed, which was filtered and washed to give the oxime (3.39 g, 20.56 mmol, 77%) as a white solid. HRMS (m/z): [M+Na]^+^ calcd. C_5_H_11_NNaO_5_ 188.0529; found, 188.0517.

To a mixture of Pt(IV)O_2_ (0.06 g) in glacial acetic acid (20 mL) was added the (D)-ribose oxime (0.60 g, 3.6 mmol). The mixture was shaken on a Parr apparatus at 45 psi hydrogen gas atmosphere for 24 h. The now clear mixture was filtered through celite and concentrated. The crude residue was dissolved in water (20 mL) and loaded onto a Dowex 50W X8 100-200 hydrogen form ion exchange column packed in water. The column was washed with water (80 mL) and then the desired ribose amine was eluted with 3N NH_4_OH solution (80 mL). The alkaline fraction was concentrated to give (2R,3S,4S)‐5‐aminopentane‐1,2,3,4‐tetrol (0.52 g, 3.4 mmol, 94%) as a light brown syrup. HRMS (m/z): [M+Na]^+^ calcd.C_5_H_13_NNaO_4_, 174.0737; found, 174.0723.

**6‐{[(2S,3S,4R)‐2,3,4,5‐Tetrahydroxypentyl]amino}‐1,2,3,4‐tetrahydropyrimidine‐2,4‐dione**

A mixture of (2R,3S,4S)‐5‐aminopentane‐1,2,3,4‐tetrol (D-ribitylamine) (0.35 g, 2.28 mmol) and 6‐chloro‐1,2,3,4‐tetrahydropyrimidine‐2,4‐dione (6-chlorouracil) (0.17g, 1.16 mmol) was heated in a pressure tube at 120°C for 3 h. The reaction was cooled to give a brown oil (0.51 g), which was used as such in the next reaction without purification. HRMS (m/z): [M+Na]^+^ calcd.C_9_H_15_N_3_NaO_6_, 284.0853; found, 284.0831.

**5‐Nitroso‐6‐{[(2S,3S,4R)‐2,3,4,5‐tetrahydroxypentyl]amino}‐1,2,3,4-tetrahydropyrimidine‐2,4‐dione**

The ribitylaminouracil-containing residue (0.22 g) was dissolved in water (10 mL) at rt and sodium nitrite (0.17g, 2.52 mmol) was added. The solution was adjusted to pH 4.6 with 3N acetic acid, stirred for 2 h, and then concentrated under vacuum. The resulting red-brown residue was dissolved in 0.15N NH_4_OH (10mL) and loaded onto a Biorad AG1-X8 resin column (formate form, 200-400 mesh). The column was washed with water (20 mL) then 0.01M formic acid (35 mL). Treatment of the column with 0.1M formic acid (400 mL) resulted in movement and discharge of a red eluate from the column, that was collected and concentrated under vacuum. The residue was recrystallised from water to give 5‐nitroso‐6‐{[(2S,3S,4R)‐2,3,4,5-tetrahydroxypentyl]amino}‐1,2,3,4-tetrahydropyrimidine‐2,4‐dione (0.17 g, 70% from (2R,3S,4S)‐5‐aminopentane‐1,2,3,4‐tetrol) as red crystals. HRMS (m/z): [M+H]^+^ calcd.C_9_H_13_N_4_O_7_, 289.0790; found, 289.0769.

**5‐amino‐6‐{[(3S,4R)‐3,4,5‐trihydroxypentyl]amino}‐1,2,3,4 tetrahydropyrimidine‐2,4‐dione (5-A-RU)**

5‐Nitroso‐6‐{[(2S,3S,4R)‐2,3,4,5-tetrahydroxypentyl]amino}‐1,2,3,4-tetrahydropyrimidine‐2,4‐dione (20.3 mg, 0.070 mmol) was suspended in milliQ water (2 mL) and heated to 80 °C under nitrogen. Sodium dithionite (40.3 mg, 0.23 mmol) was added to the red solution resulting in a colour change to pale yellow. After stirring at 80 °C for 5 min the solution was cooled in an ice bath. Analysis of this solution revealed that 5-A-RU has formed and all starting material was consumed. HRMS (m/z): [M+H]+ calcd.C_9_H_17_N_4_O_6_, 277.1143; found, 277.1126.

Under a cloud of nitrogen, the chilled solution was diluted to make a 5 mM solution. Aliquots (10 µL) were dispensed into microcentrifuge tubes, and the headspace of each tube was filled with nitrogen gas. The tubes were immediately sealed and stored at -80 °C. Before thawing an aliquot for biological testing, mass spectrometry of an aliquot was carried out to ensure that 5-A-RU had not degraded after storage.
