## Supplementary Table 1 for "Transcriptional analysis defines TCR and cytokine-stimulated MAIT cells as rapid polyfunctional effector T cells that can coordinate the immune response"

**S. Table 1:** Enriched innate signaling pathways in MAIT cells treated with *E. coli* and IL-12+IL-18 compared to 5-A-RU

| **Treatment** | **Pathways enriched in treatments vs 5-A-RU** | **Enrichment score** | **Normalized ES** | **Nominal p-value** |
| --- | --- | --- | --- | --- |
| *E. coli* | 1. Interferon alpha beta signaling | 0.827301 | 2.722886 | <0.001 |
|  | 2. Cytokine signaling in immune system | 0.590981 | 2.426092 | <0.001 |
|  | 3. Interferon signaling | 0.629474 | 2.418496 | <0.001 |
|  | 4. Interferon gamma signaling | 0.700925 | 2.311653 | <0.001 |
|  | 5. RIG-I-MDA5 mediated induction of IFN alpha beta pathways | 0.643246 | 2.25019 | <0.001 |
|  | 6. Innate immune system | 0.496261 | 2.049046 | <0.001 |
|  | 7. Negative regulators of RIG-I-MDA5 signaling | 0.684326 | 1.993789 | <0.001 |
|  | 8. TRAF6 mediated IRF7 activation | 0.646215 | 1.899533 | <0.001 |
|  | 9. TRIF mediated TLR3 signaling | 0.526647 | 1.818965 | <0.001 |
|  | 10. Activated TLR4 signaling | 0.47511 | 1.71424 | <0.001 |
|  | 11. Toll receptor cascades | 0.453692 | 1.669005 | <0.001 |
|  | 12. Nod 1/2 signaling pathway | 0.666531 | 1.904301 | 0.001645 |
|  | 13. Nucleotide binding domain leucine rich repeat containing receptor NLR signaling pathways | 0.574422 | 1.801588 | 0.001704 |
|  | 14. MYD88-MAL cascade initiated on plasma membrane | 0.469204 | 1.631594 | 0.003012 |
|  | 15. TRAF6 mediated induction of NFKB and MAP kinases upon TLR7, 8 or 9 activation | 0.488601 | 1.671944 | 0.003101 |
|  | 16. Regulation of IFN alpha signaling | 0.684114 | 1.876674 | 0.003745 |
|  | 17. NFKB and MAP kinases activation mediated by TLR4 signaling repertoire | 0.440974 | 1.498099 | 0.015924 |
|  | 18. MAP KINASE activation in TLR cascade | 0.463466 | 1.493954 | 0.024 |
| IL-12+IL-18 | 1. Interferon alpha beta signaling | 0.572527 | 1.929213 | <0.001 |
|  | 2. Interferon gamma signaling | 0.535278 | 1.785286 | <0.001 |
|  | 3. RIG-I-MDA5 mediated induction of IFN alpha beta pathways | 0.483184 | 1.695892 | <0.001 |
|  | 4. Innate immune system | 0.399903 | 1.633141 | <0.001 |
|  | 5. Interferon signaling | 0.380403 | 1.476443 | <0.001 |
|  | 6. Cytokine signaling in immune system | 0.337501 | 1.384428 | <0.001 |
|  | 7. TRAF6 mediated IRF7 activation | 0.607158 | 1.773151 | 0.001965 |
|  | 8. Negative regulators of RIG-I-MDA5 signaling | 0.608157 | 1.789948 | 0.004149 |
|  | 9. Toll receptor cascades | 0.400925 | 1.500389 | 0.004435 |
|  | 10. TRIF mediated TLR3 signaling | 0.435797 | 1.53198 | 0.011136 |
|  | 11. Activated TLR4 signaling | 0.413659 | 1.477029 | 0.015119 |
|  | 12. MYD88-MAL cascade initiated on plasma membrane | 0.403377 | 1.425398 | 0.018557 |
|  | 13. MAP KINASE activation in TLR cascade | 0.427454 | 1.386333 | 0.047109 |
