## Supplementary Figure 1 for "Transcriptional analysis defines TCR and cytokine-stimulated MAIT cells as rapid polyfunctional effector T cells that can coordinate the immune response"

### Slide 1
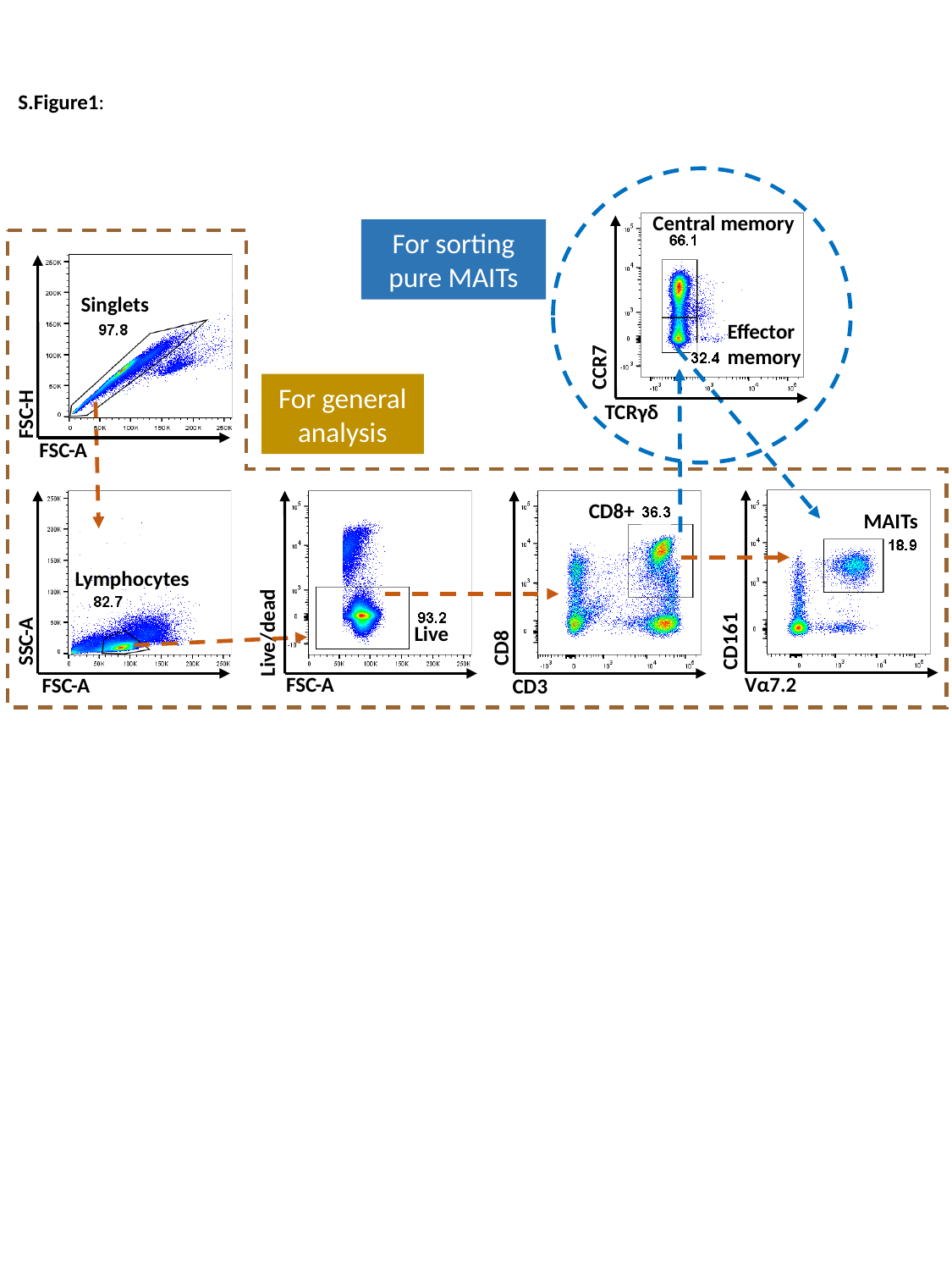

S.Figure1:
Central memory
For sorting pure MAITs
Singlets
Effector memory
CCR7
For general analysis
TCRγδ
FSC-H
FSC-A
CD8+
MAITs
Lymphocytes
Live
Live/dead
CD161
SSC-A
CD8
Vα7.2
FSC-A
FSC-A
CD3
