## Supplementary Figure 2 for "Transcriptional analysis defines TCR and cytokine-stimulated MAIT cells as rapid polyfunctional effector T cells that can coordinate the immune response"

### Slide 1
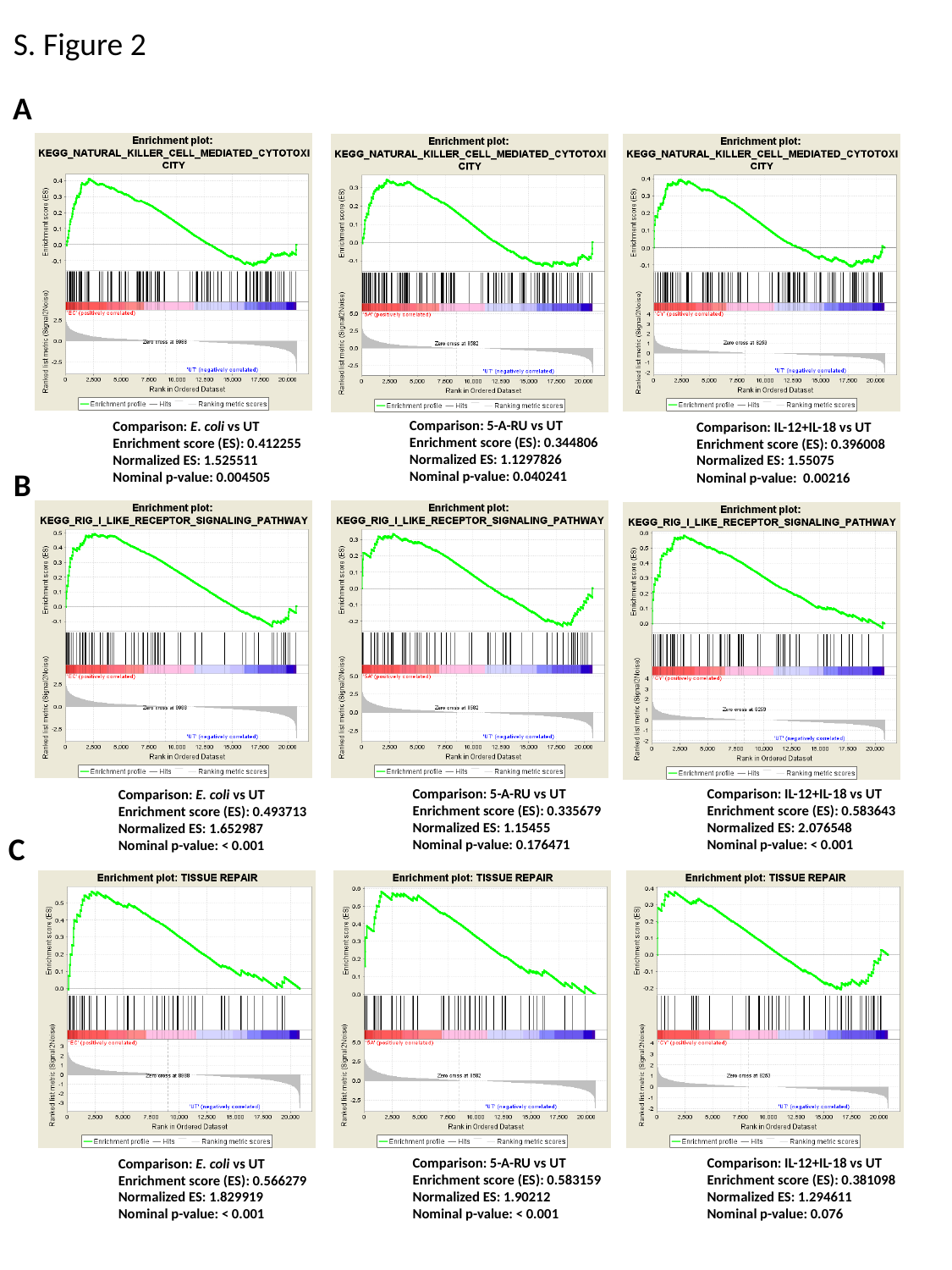

S. Figure 2
A
Comparison: 5-A-RU vs UT
Enrichment score (ES): 0.344806
Normalized ES: 1.1297826
Nominal p-value: 0.040241
Comparison: E. coli vs UT
Enrichment score (ES): 0.412255
Normalized ES: 1.525511
Nominal p-value: 0.004505
Comparison: IL-12+IL-18 vs UT
Enrichment score (ES): 0.396008
Normalized ES: 1.55075
Nominal p-value: 0.00216
B
Comparison: 5-A-RU vs UT
Enrichment score (ES): 0.335679
Normalized ES: 1.15455
Nominal p-value: 0.176471
Comparison: IL-12+IL-18 vs UT
Enrichment score (ES): 0.583643
Normalized ES: 2.076548
Nominal p-value: < 0.001
Comparison: E. coli vs UT
Enrichment score (ES): 0.493713
Normalized ES: 1.652987
Nominal p-value: < 0.001
C
Comparison: 5-A-RU vs UT
Enrichment score (ES): 0.583159
Normalized ES: 1.90212
Nominal p-value: < 0.001
Comparison: IL-12+IL-18 vs UT
Enrichment score (ES): 0.381098
Normalized ES: 1.294611
Nominal p-value: 0.076
Comparison: E. coli vs UT
Enrichment score (ES): 0.566279
Normalized ES: 1.829919
Nominal p-value: < 0.001
