## Supplementary figures and images for "Transcriptional analysis defines TCR and cytokine-stimulated MAIT cells as rapid polyfunctional effector T cells that can coordinate the immune response"

### Supplementary Figure 3

#### Slide 1
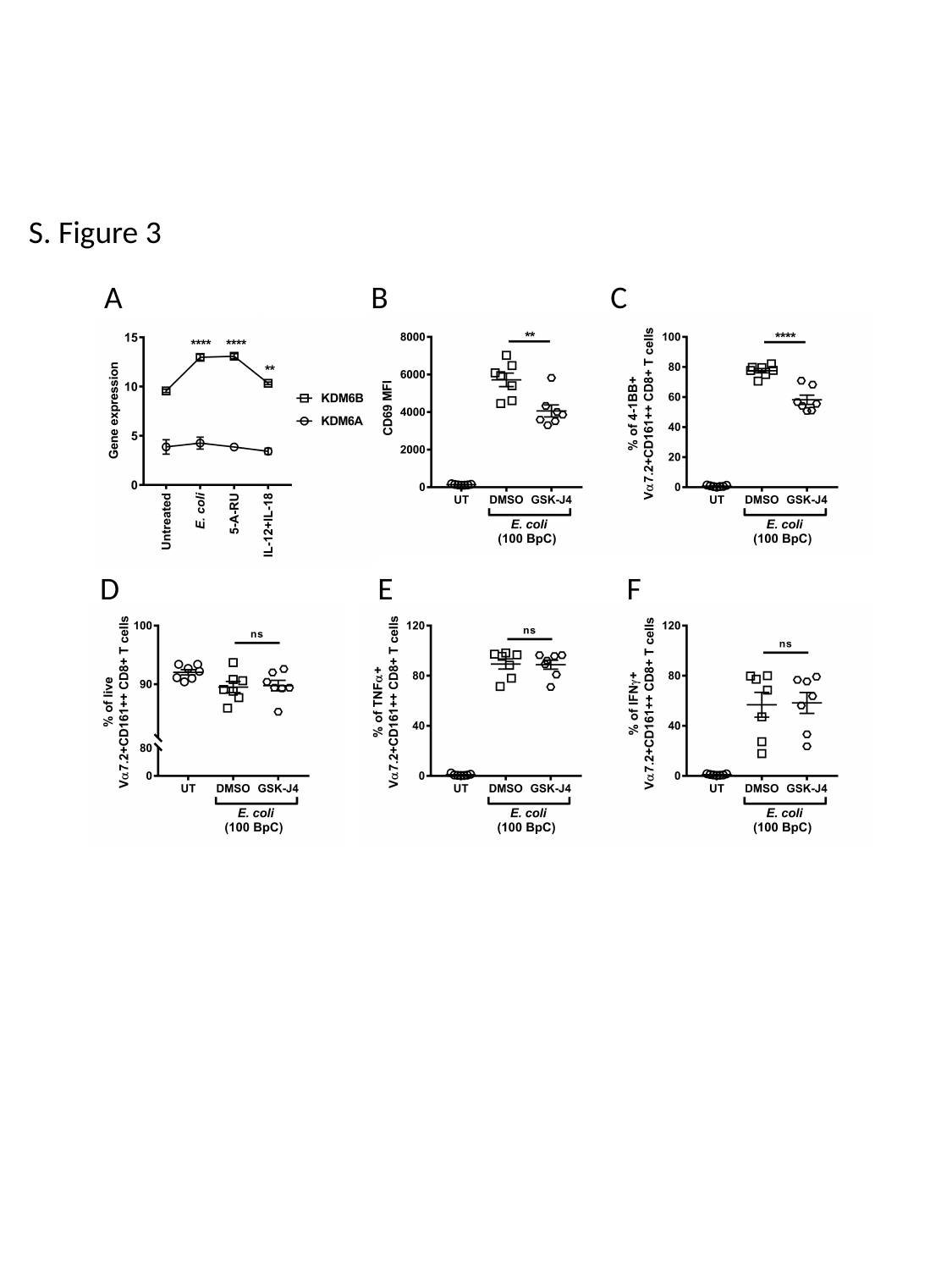

S. Figure 3
A
B
C
D
E
F

### Supplementary Figure 4

#### Slide 1
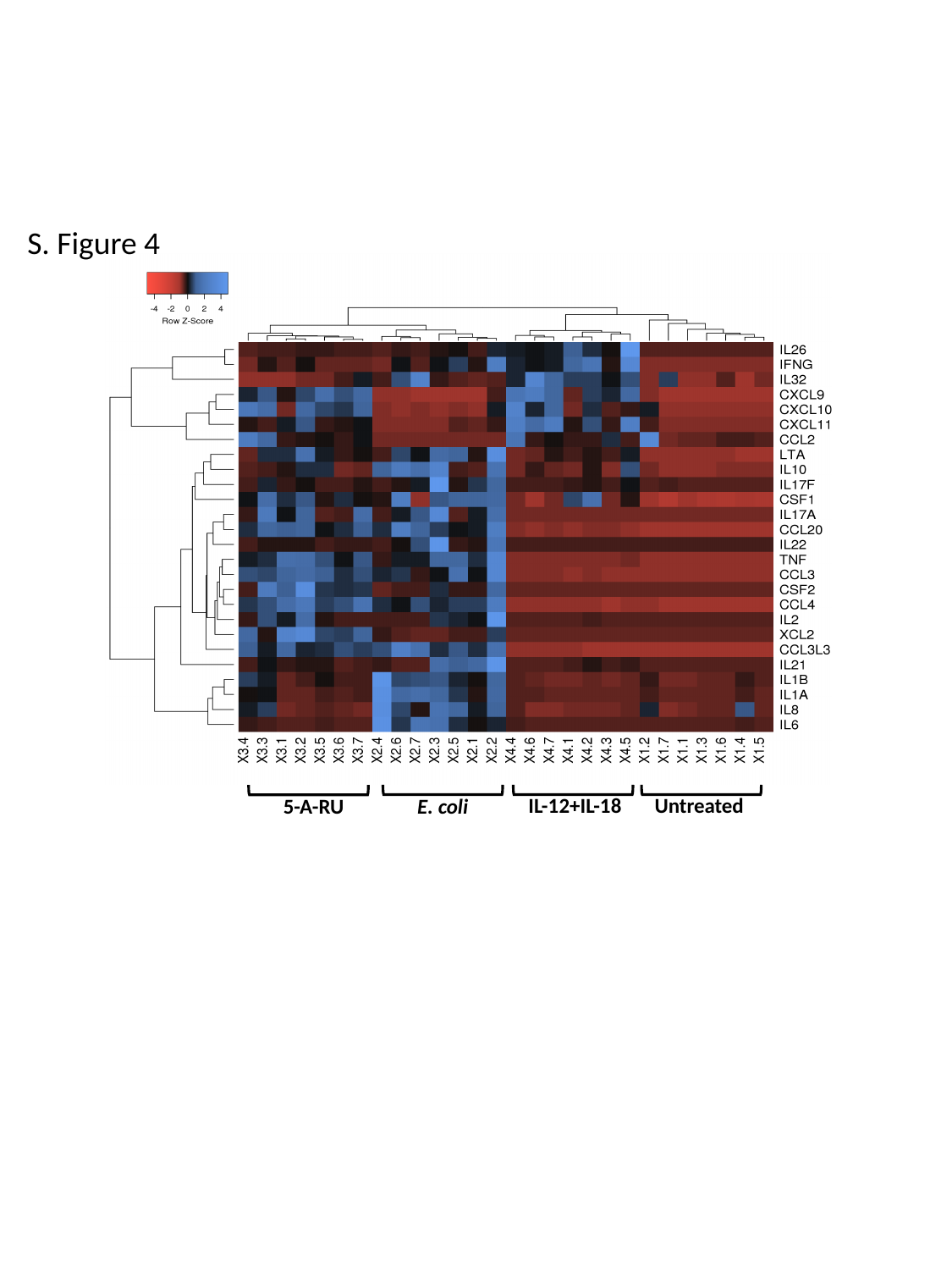

S. Figure 4
IL-12+IL-18
Untreated
5-A-RU
E. coli

### Supplementary Figure 5

#### Slide 1
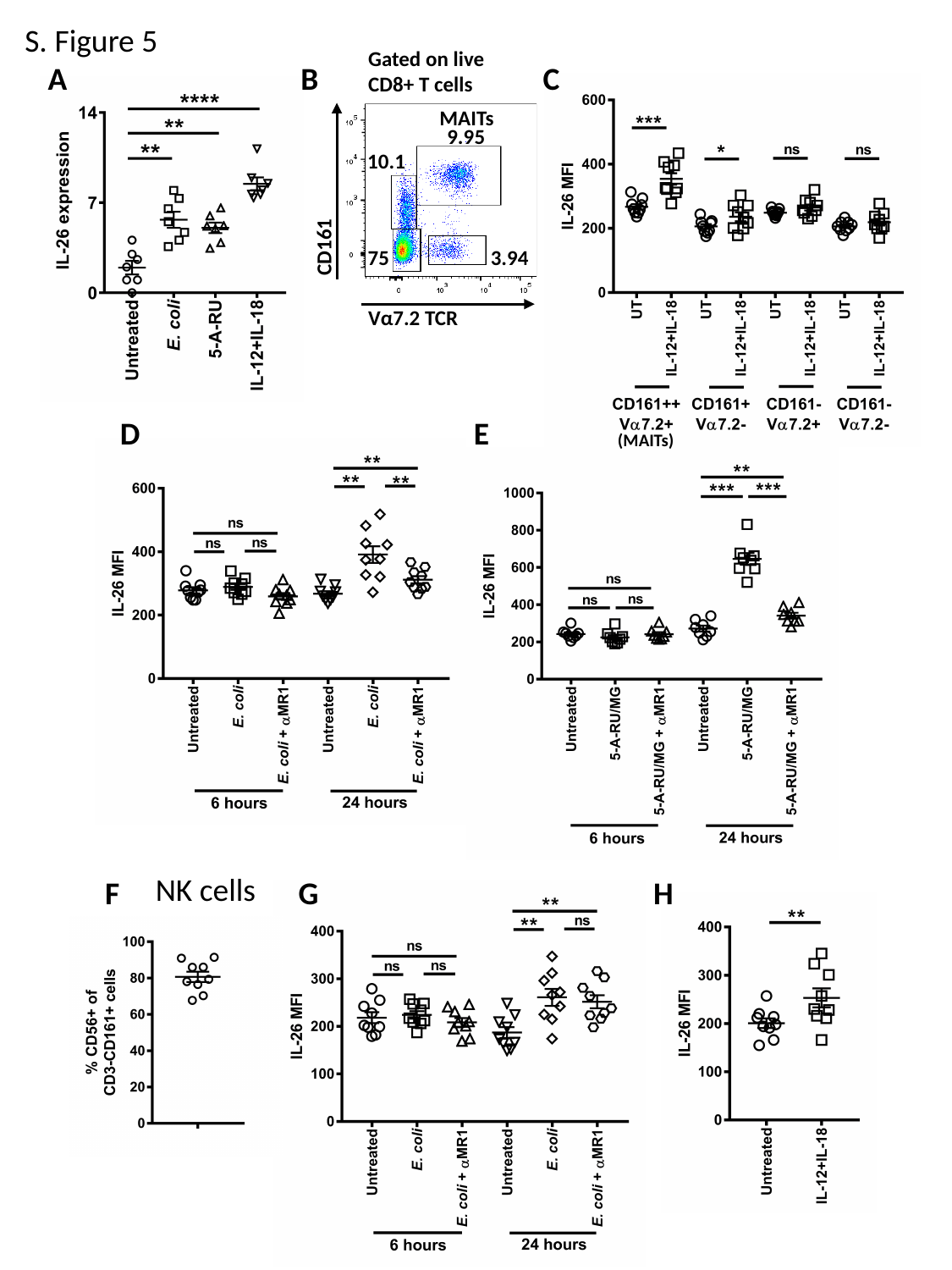

S. Figure 5
Gated on live
CD8+ T cells
A
B
C
9.95
10.1
75
3.94
MAITs
CD161
Vα7.2 TCR
D
E
(MAITs)
NK cells
F
G
H

### Supplementary Figure 6

#### Slide 1
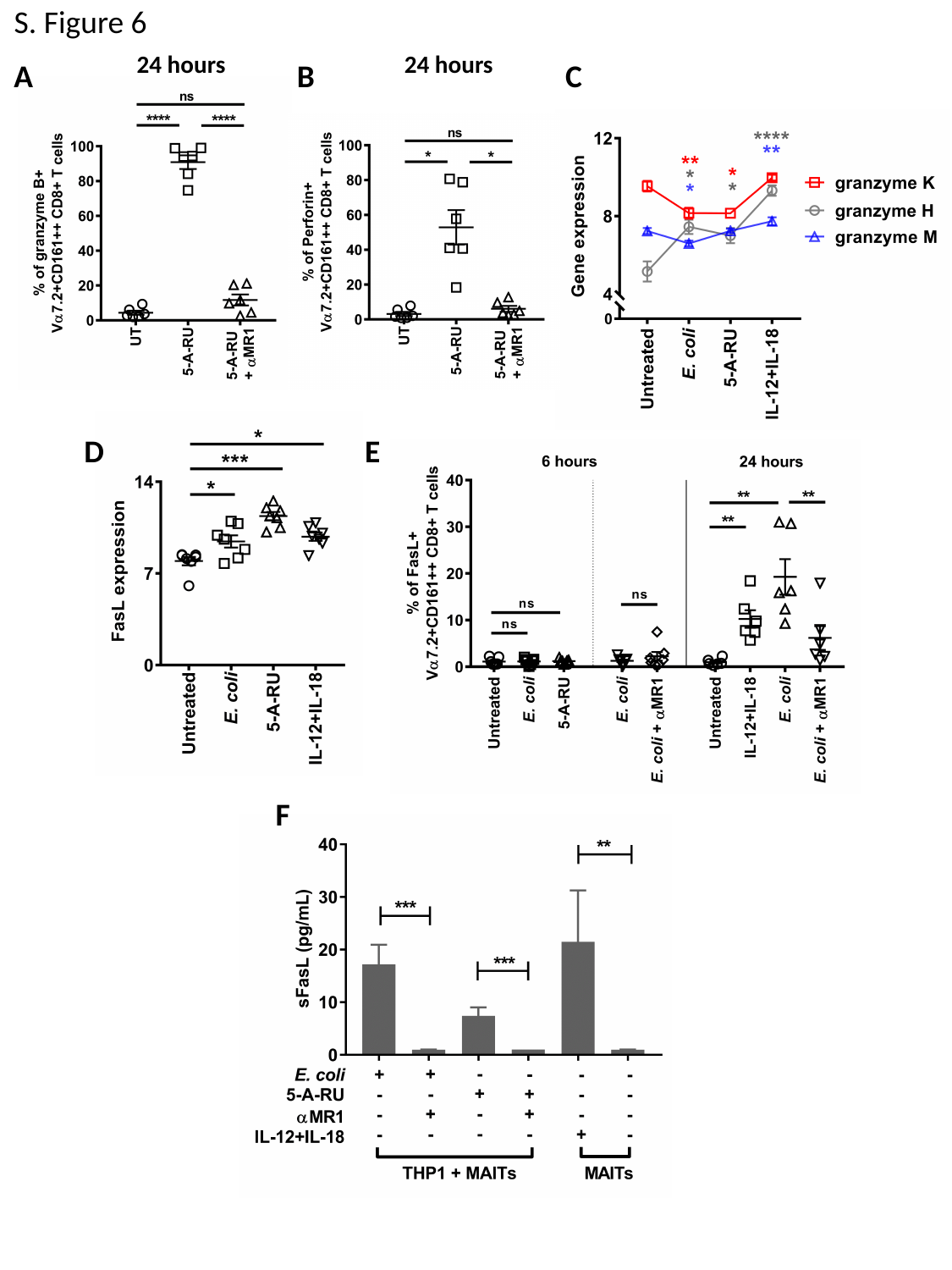

S. Figure 6
24 hours
24 hours
A
B
C
E
D
F

### Supplementary Figure 7

#### Slide 1
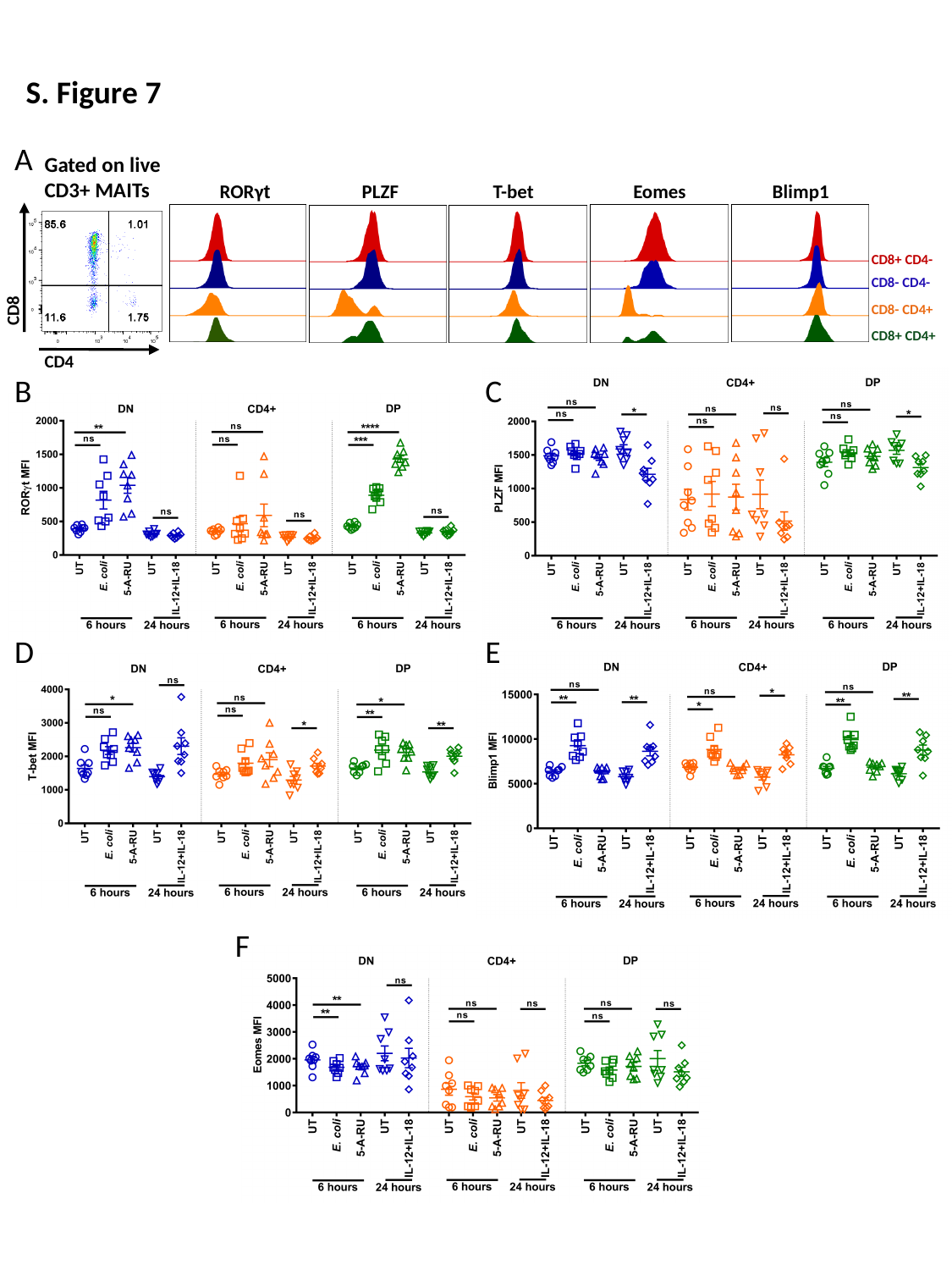

S. Figure 7
A
Gated on live
CD3+ MAITs
RORγt
PLZF
T-bet
Eomes
Blimp1
CD8+ CD4-
CD8- CD4-
CD8
CD8- CD4+
CD8+ CD4+
CD4
B
C
D
E
F
